# Self-assembly of the RZZ complex into filaments drives kinetochore expansion in the absence of microtubule attachment

**DOI:** 10.1101/282707

**Authors:** Cláudia Pereira, Rita M. Reis, José B. Gama, Dhanya K. Cheerambathur, Ana X. Carvalho, Reto Gassmann

## Abstract

The kinetochore is a dynamic multi-protein assembly that forms on each sister chromatid and interacts with microtubules of the mitotic spindle to drive chromosome segregation. In animals, kinetochores without attached microtubules expand their outermost layer into crescent and ring shapes to promote microtubule capture and spindle assembly checkpoint (SAC) signalling. Kinetochore expansion is an example of protein co-polymerization, but the mechanism is not understood. Here, we present evidence that kinetochore expansion is driven by oligomerization of the Rod-Zw10-Zwilch (RZZ) complex, an outer kinetochore component that recruits the motor dynein and the SAC proteins Mad1-Mad2. Depletion of ROD in human cells suppresses kinetochore expansion, as does depletion of Spindly, the adaptor that connects RZZ to dynein, while dynein itself is dispensable. Expansion is also suppressed by mutating ZWILCH residues implicated in Spindly binding. Conversely, supplying cells with excess ROD facilitates kinetochore expansion under otherwise prohibitive conditions. Using the *C. elegans* early embryo, we demonstrate that ROD-1 has a concentration-dependent propensity for oligomerizing into *µ*m-scale filaments, and we identify the ROD-1 β-propeller as a key regulator of self-assembly. Finally, we show that a minimal ROD-1-Zw10 complex efficiently oligomerizes into filaments *in vitro*. Our results suggest that RZZ’s capacity for oligomerization is harnessed by kinetochores to assemble the expanded outermost domain, in which RZZ filaments serve as recruitment platforms for SAC components and microtubule-binding proteins. Thus, we propose that RZZ self-assembly into filaments underlies the adaptive change in kinetochore size that contributes to chromosome segregation fidelity.

## INTRODUCTION

Equal segregation of chromosomes to daughter cells during cell division requires interactions between spindle microtubules and the kinetochore, a complex multi-protein assembly that forms at the centromeric locus of each sister chromatid. In animal cells, kinetochores only gain access to microtubules once the nuclear envelope breaks down. For timely bi-orientation of sister chromatids on the mitotic spindle, kinetochores must be efficient at capturing microtubules and converting initial lateral microtubule contacts to the stable end-on attachments that drive chromosome segregation [1]. As a safeguard, the spindle assembly checkpoint (SAC) generates an inhibitory “wait anaphase” signal at kinetochores that have not yet established stable attachments to microtubules, thereby ensuring that the cell proceeds with anaphase and mitotic exit only once all sister kinetochores have bi-oriented [2]. Elucidating the molecular mechanisms that promote microtubule capture and SAC signalling at kinetochores remains an important challenge.

Kinetochores assemble in a largely hierarchical manner starting from CENP-A-containing nucleosomes and the associated protein CENP-C, which is part of the 16-subunit constitutive centromere-associated network (CCAN; also referred to as the inner kinetochore) [3]. The CCAN then directs the assembly of the microtubule interface (or outer kinetochore), consisting of the 10-subunit Knl1, Mis12 complex, Ndc80 complex (KMN) network. The KMN network mediates end-on microtubule attachment through the Ndc80 complex and provides a platform for generation of the mitotic checkpoint complex (MCC), consisting of BubR1 (also known as Mad3), Bub3, Mad2, and Cdc20, which prevents mitotic exit by inhibiting the E3 ubiquitin ligase anaphase promoting complex/cyclosome (APC/C) [4]. Knl1 acts as the scaffold for the recruitment of Bub3, BubR1, and the kinase Bub1 [5], which is thought to promote MCC formation by directly binding to a Mad1-Mad2 template that catalyzes MCC assembly [6–10]. Knl1 and Bub1 are also implicated in the recruitment of the three-subunit Rod-Zw10-Zwilch (RZZ) complex [11–15], which is one of several components that localize peripherally to the KMN network and include the microtubule-associated protein (MAP) CENP-F and the microtubule motors dynein and CENP-E. RZZ recruits dynein and its co-factor dynactin through a direct interaction with the dynein adaptor Spindly [15,16]. RZZ is also required for Mad1-Mad2 recruitment, and recent work suggests that RZZ’s contribution to SAC signalling may be distinct from that of Bub1 [13,17].

Kinetochores have long been known to respond to the absence of microtubules by expanding into ring and crescent shapes that encircle the centromere [18,19]. Kinetochore expansion occurs through addition of new material rather than re-distribution of existing protein and is characterized by the formation of an outermost domain, called the fibrous corona based on its appearance in electron microscopy [18,20]. Kinetochore expansion is particularly prominent when mitosis with unattached kinetochores is prolonged by treatment with microtubule depolymerizing drugs [20,21], but kinetochores also expand in unperturbed cells in early prometaphase, when kinetochores are devoid of microtubules [22]. Kinetochore expansion is proposed to make two critical contributions to the fidelity of chromosome segregation: it increases the amount of kinetochore-localized SAC proteins, including Mad1 and Mad2, implying that more MCC can be generated per unattached kinetochore, and it distributes MAPs such as CENP-F, CENP-E and dynein over a larger surface, which facilitates lateral microtubule capture [19,21–24]. Consequently, kinetochore expansion is predicted to both expedite the formation of end-coupled kinetochore-microtubule attachments and amplify the checkpoint signal [21,22], which may be particularly relevant when anaphase has to be delayed in the presence of just a few or even a single unattached kinetochore [25].

Mechanistically, kinetochore expansion remains poorly understood but is thought to be an example of multiprotein co-polymerization [21]. Recent work using super-resolution light microscopy in *Xenopus* egg extracts identified an expandable kinetochore module consisting of *µ*m-long fibers that grow out from centromeric chromatin along chromosome arms. Fibrous extensions emanating from mitotic chromosomes have also been observed in *C. elegans* embryos treated with nocodazole [26], and filaments containing kinetochore components surround chromosomes in the *C. elegans* meiosis I embryo [27,28]. Intriguingly, recent analysis of reconstituted human RZZ by cryo-electron microscopy confirmed an earlier prediction that the ROD subunit is structurally related to membrane coat proteins such as Clathrin and subunits of the COPI and COPII complexes [16,29]. The underlying common design, which consists of an N-terminal β-propeller domain and C-terminally located α-solenoid motifs, enables coat proteins to form higher-order assemblies around vesicles that act as scaffolds to direct membrane traffic [30,31].

Here, using cultured human cells, the *C. elegans* early embryo, and purified proteins, we demonstrate that RZZ is capable of oligomerizing into *µ*m-scale filaments and present evidence that Rod is the critical subunit for self-assembly, as predicted by its architectural similarity with membrane coat proteins. Our results suggest that RZZ’s propensity for oligomerization is harnessed at kinetochores to drive the assembly of the expanded outer domain, in which RZZ filaments serve as platforms for the recruitment of SAC proteins and MAPs.

## RESULTS

### Kinetochore expansion requires the RZZ complex and Spindly but not dynein-dynactin

To examine the role of the kinetochore dynein module (RZZ-Spindly-dynein-dynactin) in kinetochore expansion, we incubated HeLa cells with nocodazole to depolymerize microtubules and used immunostaining for the outer kinetochore proteins CENP-E and CENP-F to assess crescent formation (Fig. 1A). In cells treated with control siRNA, CENP-E and CENP-F expanded into crescents that partially encircled the compact inner kinetochore, marked by CENP-C, as expected (Fig. 1B). Depletion of the RZZ subunit ROD by RNAi, which eliminated Spindly localization to kinetochores, supported CENP-E and CENP-F recruitment, but kinetochores no longer expanded into crescents (Fig. 1B). Measurements of kinetochore fluorescence confirmed that ROD depletion reduced both the volume occupied by CENP-E and CENP-F and their overall levels (Fig. 1C - F). Depletion of Spindly also reduced kinetochore expansion, albeit not to the same degree as depletion of ROD (Fig. 1B - F). By contrast, depletion of the dynactin subunit DCTN1, which prevents kinetochore recruitment of both dynactin and dynein, did not affect kinetochore expansion, as judged by immunostaining for Spindly (Fig. 1G - I). We conclude that kinetochore expansion requires RZZ and Spindly but is independent of dynein-dynactin.

**Figure 1:**
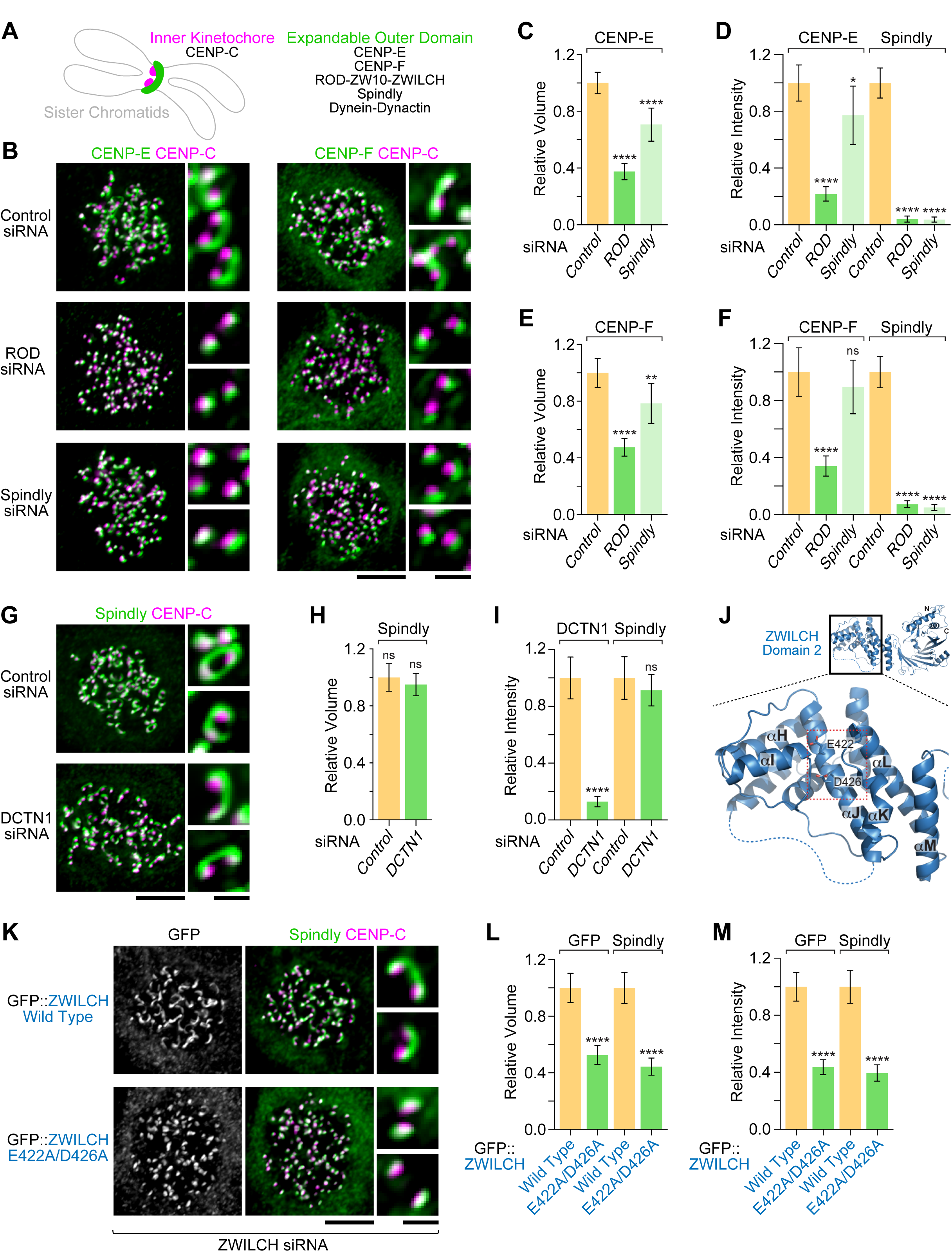
Kinetochore expansion requires the RZZ complex and Spindly but is independent of dynein-dynactin. **(A)** Cartoon showing the crescent shape characteristic of the expanded outer kinetochore, which encircles the compact inner kinetochore. Components analyzed in this figure are listed on the right. **(B)** Immunofluorescence images showing that kinetochore expansion in nocodazole is inhibited after RNAi-mediated depletion of ROD or Spindly. Scale bar, 5 *µ*m; blow-ups, 1 *µ*m. **(C)** - **(F)** Quantification of relative kinetochore volume *(C, E)* and signal intensity *(D, F)* for the components indicated on top, based on fluorescence measurements in images as shown in *(B)*. For each condition, the mean value per kinetochore was determined for individual cells. Final values are shown as the mean of mean (n = 20 cells), normalized to the control. Error bars denote the 95% confidence interval. Statistical significance was determined by one-way ANOVA followed by Bonferroni’s multiple comparison test. \*\*\*\**P* < 0.0001; \*\**P* < 0.01; \**P* < 0.05; ns = not significant, *P* > 0.05. **(G)** Immunofluorescence images showing that depletion of the dynactin subunit DCTN1 does not affect kinetochore expansion. Scale bar, 5 *µ*m; blow-ups, 1 *µ*m. **(H), (I)** Quantification of relative kinetochore volume *(H)* and signal intensity *(I)*, determined and plotted as described for *(C) - (F)*. The t-test was used to determine statistical significance. \*\*\*\**P* < 0.0001; ns = not significant, *P* > 0.05. **(J)** Cartoon model of ZWILCH (PDB ID 3IF8) obtained with PyMOL. Dotted lines mark loop regions that were not visible in the crystal structure and were added to improve clarity. The red boxed region indicates the position of E422 and D426. **(K)** Immunofluorescence images showing that the ZWILCH mutant E422A/D426A does not support kinetochore expansion. Scale bar, 5 *µ*m; blow-ups, 1 *µ*m. **(L), (M)** Quantification of relative kinetochore volume *(L)* and signal intensity *(M)*, determined and plotted as described for *(C) - (F)*. The t-test was used to determine statistical significance. \*\*\*\**P* < 0.0001.

### ZWILCH residues implicated in Spindly binding are required for kinetochore expansion

Human Spindly directly interacts with the ROD β-propeller [16], and the β-propeller is essential for Spindly recruitment to kinetochores [15]. The atomic-resolution structure of the RZZ subunit ZWILCH, which forms a tight complex with the ROD β-propeller, features a conserved surface-exposed patch of unknown function in domain 2 [29] (Fig. 1J). In *C. elegans*, two acidic residues in this patch are required for an interaction between ZWL-1^Zwilch^ and SPDL-1^Spindly^[15]. We used the Flp-In system in HeLa cells and RNAi-based molecular replacement to assess the consequences of introducing analogous mutations (E422A/D426A) into human ZWILCH (Fig. 1J). Expression of transgene-encoded GFP::ZWILCH, harboring silent mutations that confer RNAi resistance, supported recruitment of Spindly to kinetochores and facilitated kinetochore expansion in nocodazole, as judged by immunostaining for GFP and Spindly (Fig. 1 K - M). By contrast, GFP::ZWILCH(E422A/D426A), while supporting Spindly recruitment to kinetochores, inhibited kinetochore expansion to a similar degree as depletion of ROD (Fig. 1 K - M). We conclude that kinetochore expansion requires two conserved acidic ZWLICH residues that in *C. elegans* ZWL-1^Zwilch^ are implicated in an interaction with SPDL-1^Spindly^.

### The expanded outer kinetochore is a distinct multi-protein domain that can be dissociated from the KMN network

We found that, following a 4-h incubation in nocodazole, a 30-min treatment of mitotic cells with the CDK1 inhibitor RO-3306 resulted in complete detachment of the expanded outer domain from the centromere (Fig. 2A). Immunostaining showed that crescent-like assemblies in the cytoplasm, separated by several *µ*m from the closest sister kinetochore pair, marked by CENP-C or human anti-centromere antibodies (ACA), contained ZW10, ZWILCH, Spindly, the dynactin subunit DCTN1, CENP-E, MAD1, and MAD2 (Fig. 2B, D). By contrast, we could not detect KNL1, BUB1, BUBR1, BUB3, CENP-F, HEC1, DSN1, or CENP-C on detached crescents (Fig. 2D; Fig. S1). Consistently, detached crescents were never observed after depletion of ROD or Spindly. We conclude that kinetochore expansion involves the assembly of a distinct multi-protein domain with defined composition that is separable from the inner kinetochore and the KMN network.

**Figure 2:**
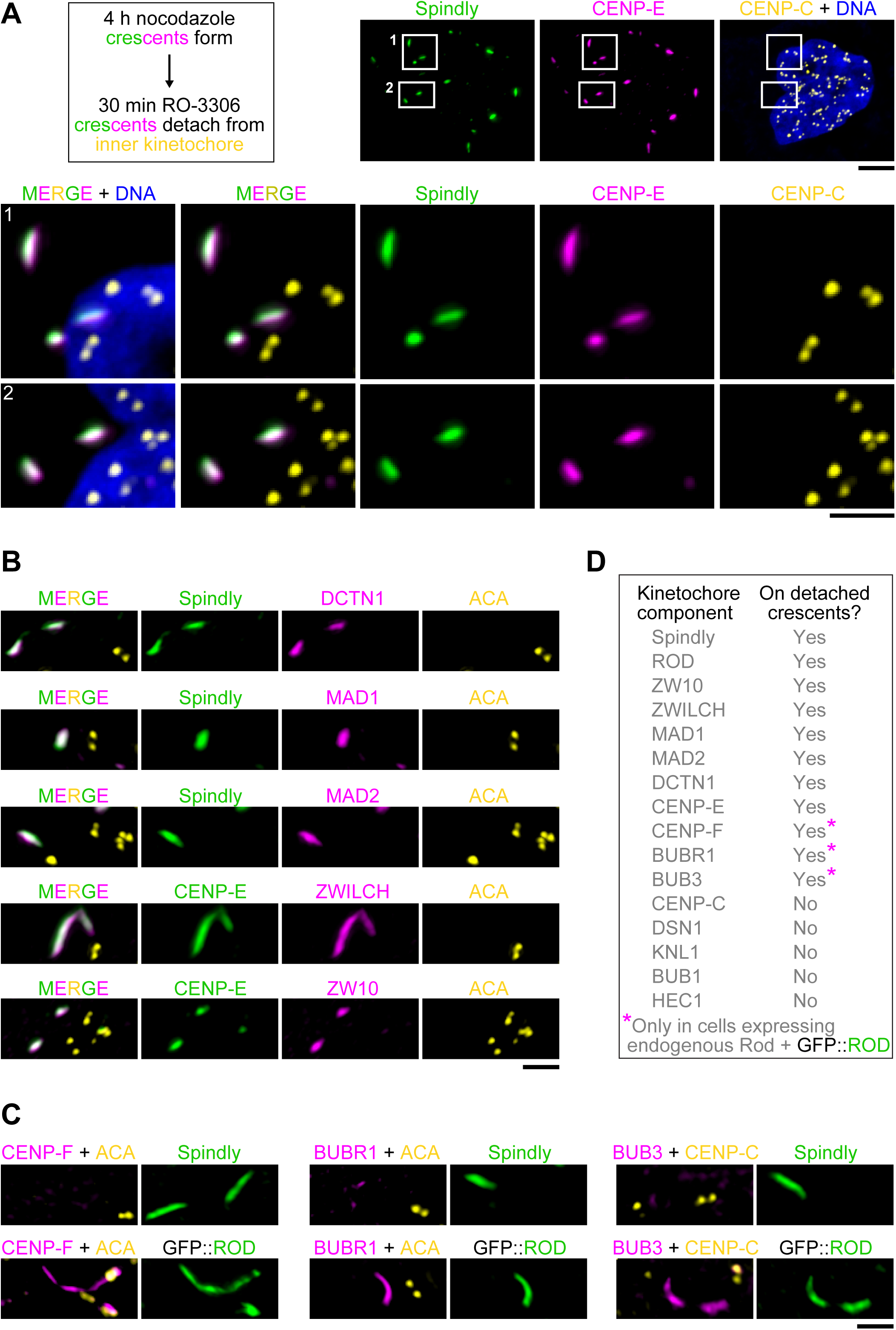
The expanded outer kinetochore is a distinct multi-protein domain that can be dissociated from the KMN network. **(A)** *(left)* Summary of the experimental regime to detach the expanded kinetochore domain (crescents) from the centromere. *(right)*, *(bottom)* Immunofluorescence images showing crescents, marked by CENP-E and Spindly, that are fully detached from the inner kinetochore, marked by CENP-C. Boxed regions are shown separately at higher magnification. Scale bar, 5 *µ*m; blow-ups, 2 *µ*m. **(B)** Immunofluorescence images showing the composition of detached crescents (see also Fig. S1). Scale bar, 2 *µ*m. **(C)** Immunofluorescence images highlighting the components that only localize to detached crescents in cells expressing exogenous GFP::ROD. Scale bar, 2 *µ*m. **(D)** Table summarizing the protein composition of detached crescents, based on the immunofluorescence analysis shown in *(A)* - *(C)* and in Fig. S1.

The robust presence of MAD1-MAD2 on detached kinetochore crescents in the absence of detectable KNL1 or BUB1 implied that RZZ is directly involved in MAD1-MAD2 recruitment. Interestingly, when we generated detached crescents in cells expressing transgene-encoded GFP::ROD on top of endogenous ROD, CENP-F, BUBR1, and BUB3 were consistently detected on detached crescents (Fig. 2C, D). Thus, over-expression of GFP::ROD recruited outer kinetochore components that were normally not present at detached crescents. Of note, prior work showed that CENP-F has affinity for recombinant GFP::RZZ [16], suggesting that RZZ may promote CENP-F recruitment though a direct interaction. Regarding the SAC, these results imply that RZZ is capable of recruiting MCC components to the expanded outer kinetochore independently of KNL1-BUB1.

### Expression of exogenous ROD promotes kinetochore expansion

To test whether over-expression of GFP::ROD promotes kinetochore expansion *per se*, we expressed GFP::ROD under conditions that usually compromise expansion. Consistent with previous reports [14,17], RNAi-mediated depletion of KNL1 reduced the amount of endogenous RZZ recruited to kinetochores, as revealed by immunostaining for ZWILCH, and kinetochore expansion was significantly suppressed (Fig. 3A). In this and the following experiments, the absence of BUB1 signal at kinetochores was used to confirm efficient depletion of KNL1 (Fig. 3A). Strikingly, expression of GFP::ROD in KNL1-depleted cells produced robustly expanded kinetochores (Fig. 3B). Indeed, GFP::ROD-containing crescents in KNL1-depleted cells were often larger than GFP::ROD crescents in cells treated with control siRNA and occasionally connected kinetochores of adjacent chromosomes (Fig. 3C). GFP::ROD-containing crescents and rings that were partially or even fully detached from sister kinetochore pairs were also prevalent in KNL1-depleted cells but were never observed in cells treated with control siRNA (Fig. 3B, C). The protein composition of detached GFP::ROD crescents was identical to the detached crescents generated by acute CDK1 inhibition (Fig. 3C and data not shown). As expected, co-depletion of KNL1 and Spindly suppressed kinetochore expansion in cells expressing GFP::ROD (Fig. 3D). We also examined the effect of a GFP::ROD mutant missing its N-terminal β-propeller domain (Δ1-375), which we previously showed localizes to kinetochores but prevents recruitment of ZWILCH and Spindly [15]. GFP::ROD(Δ1-375) did not support expansion, as we pointed out previously [15], and depletion of KNL1 greatly reduced the kinetochore signal of the ROD mutant compared to cells treated with control siRNA (Fig. S2).

**Figure 3:**
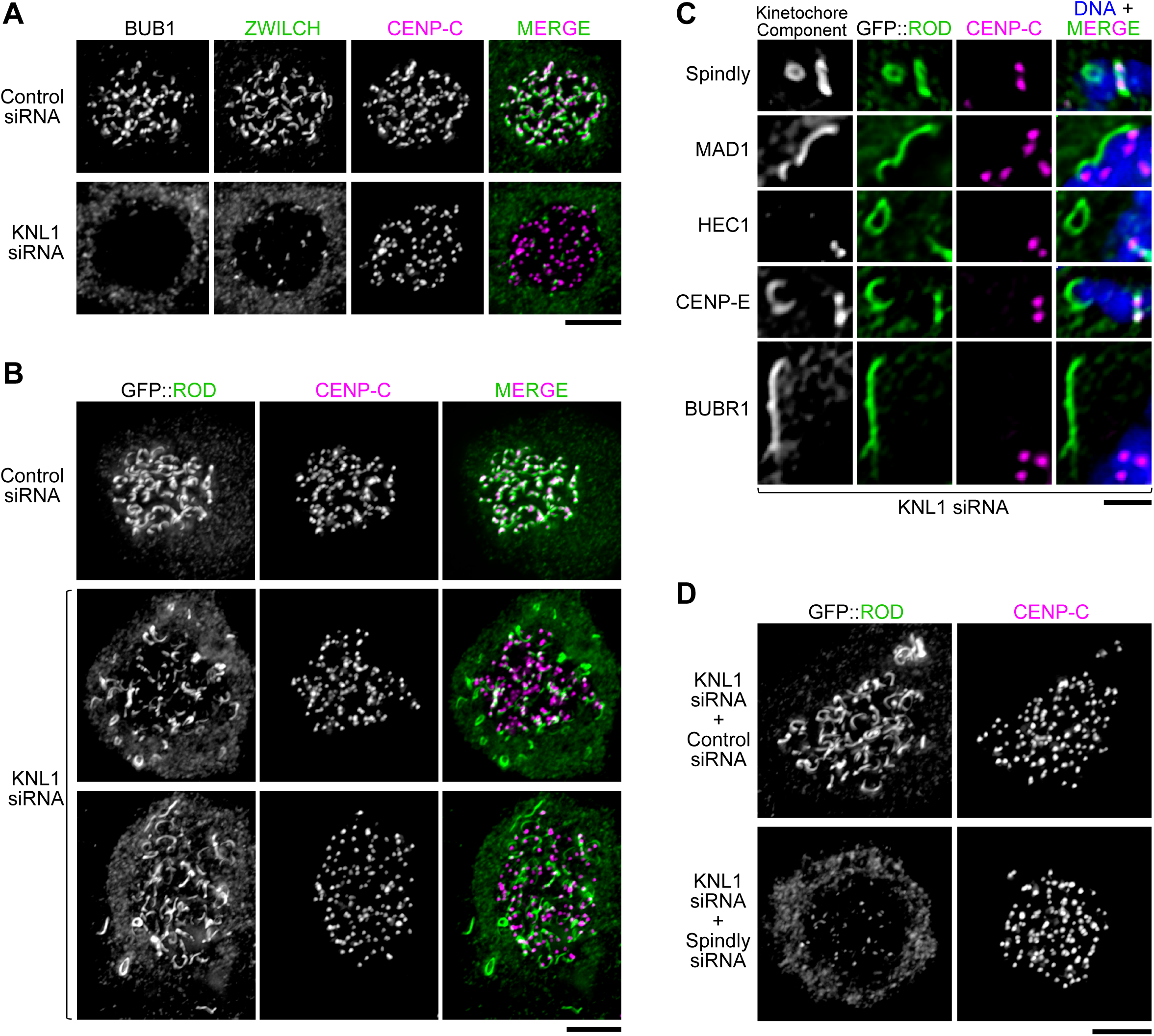
Expression of exogenous GFP::ROD promotes kinetochore expansion. **(A)** Immunofluorescence images showing that KNL1 depletion reduces RZZ levels at kinetochores and prevents kinetochore expansion. BUB1 staining serves as a readout for KNL1 depletion. Scale bar, 5 *µ*m. **(B)** Immunofluorescence images showing that exogenous expression of GFP::ROD promotes kinetochore expansion in KNL1-depleted cells. As in *(A)*, efficient KNL1 depletion was confirmed by co-staining the same cells for BUB1. Note that the expanded domains are only loosely associated with the centromere, marked by CENP-C. Scale bar, 5 *µ*m. **(C)** Examples of detached crescents and rings in GFP::ROD-expressing cells depleted of KNL1, showing co-localization with other kinetochore components. Scale bar, 2 *µ*m. **(D)** Immunofluorescence images showing that GFP::ROD is unable to support kinetochore expansion in the absence of Spindly. Scale bar, 5 *µ*m.

Taken together, these experiments suggest that increased ROD levels facilitate robust kinetochore expansion even when kinetochores have a reduced capacity for RZZ recruitment, and that this depends on Spindly and the ROD β-propeller. This is consistent with an expansion mechanism based on self-assembly of RZZ, for which we present direct evidence below. The GFP::ROD(Δ1-375) signal likely represents the RZZ pool directly recruited by KNL1-BUB1 without expansion, while the GFP::ROD signal also reflects higher-order assemblies of RZZ. These experiments also show that KNL1-BUB1 are required to tether the expanded outer domain to the core kinetochore.

### *C. elegans* ROD-1 is capable of self-assembly into *µ*m-scale filaments *in vivo*

Rod proteins share a common architecture with membrane coat precursors, which self-assemble into higher-order structures [16]. Self-assembly of ROD into filaments could underlie the central role of ROD in kinetochore expansion that we describe above, but there is currently no evidence that Rod proteins can form oligomeric assemblies. In our search for such evidence, we turned to the nematode *C. elegans*. In mature oocytes and meiosis I embryos, small filamentous assemblies, referred to as ‘linear elements’, are present in the cytoplasm and prominently encircle the bivalent chromosomes [27,28] (Fig. S3). Linear elements contain RZZ and other kinetochore proteins, including KNL-1, BUB-1, SPDL-1^Spindly^, and dynein [27,28] (Fig. S3). Depletion of ROD-1 prevented the formation of linear elements (Fig. S3A), suggesting that they may represent oligomeric assemblies of ROD-1. Intriguingly, we found that after tagging endogenous ROD-1 with GFP using CRISPR/Cas9-mediated genome editing, GFP::ROD-1 filaments were not only present in meiosis but also appeared in the cytoplasm of early multicellular embryos (Fig. 4A). Live imaging revealed that *µ*m-scale GFP::ROD-1 filaments formed with highly reproducible kinetics during mitosis at the 8-cell stage, but not in earlier divisions (Fig. 4A; Movie S1). In addition to forming filaments, GFP::ROD-1 localized transiently to holocentric kinetochores and was enriched in nuclei in early embryos at all stages (Fig. 4A). At the 8-cell stage, filaments formed rapidly (within ∼40 s) in nuclei about 3 min prior to NEBD, at about the same time when the GFP::ROD-1 signal first appeared on kinetochores (Fig. 4B; Movie S2). GFP::ROD-1 filaments remained distinct from mitotic chromosomes and segregated to daughter cells by clustering at spindle poles (Fig. 4B; Fig. S4A; Movie S2). At the end of mitosis, the filaments largely disassembled before forming again in nuclei prior to NEBD of the subsequent division (Fig. S4B; Movie S2). Interestingly, depletion of SPDL-1^Spindly^ had no effect on the formation of GFP::ROD-1 filaments but prevented clustering at spindle poles (Fig. S4B), suggesting that the filaments become tethered to spindle poles via dynein.

**Figure 4:**
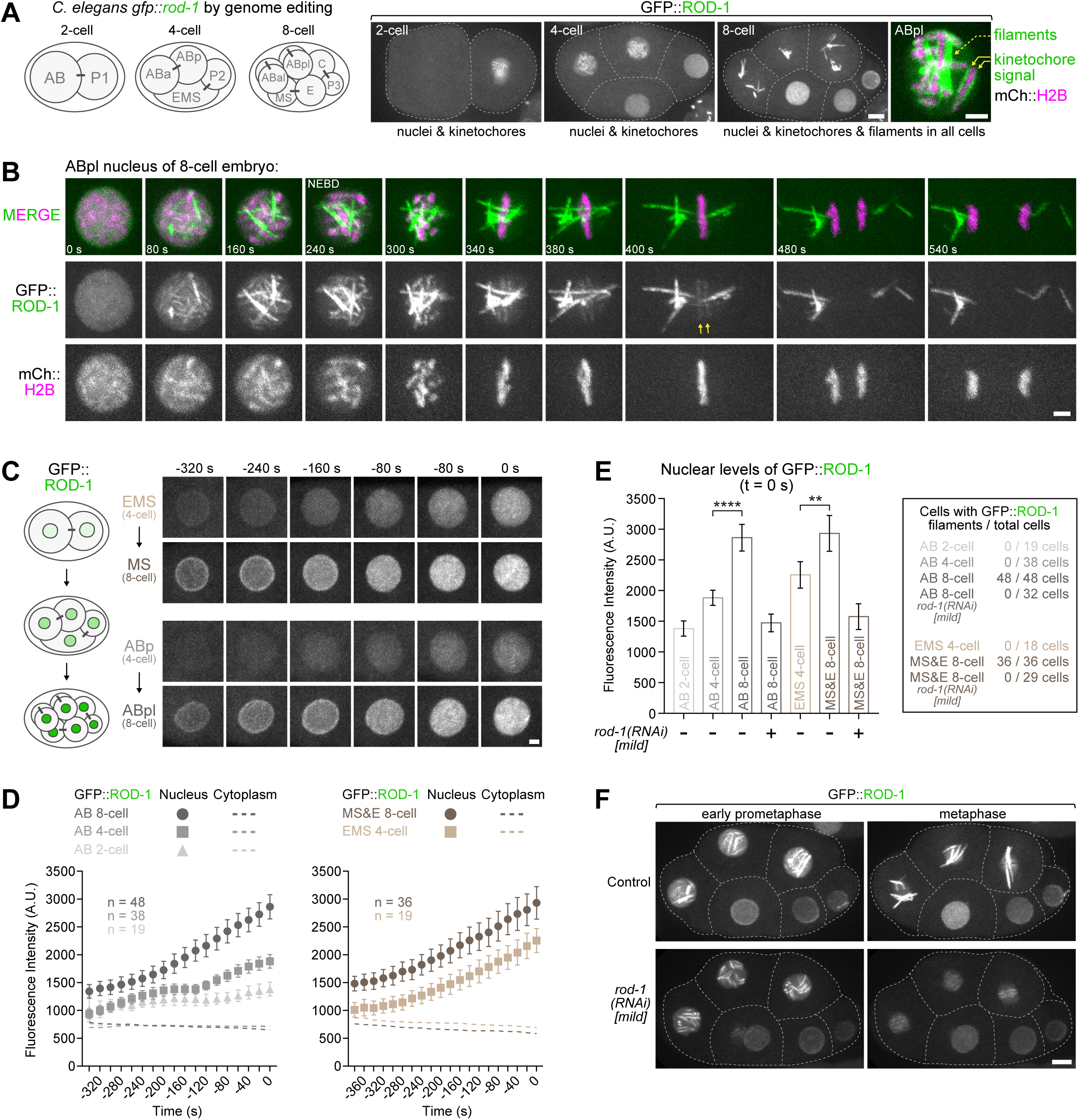
*C. elegans* ROD-1 is capable of self-assembly into *µ*m-scale filaments *in vivo*. **(A)** *(left)* Schematic of the *C. elegans* early embryo at the 2-, 4-, and 8-cell stage. Names of individual cells are indicated. Bars connecting cells indicate the they originated from the same mother cell. *(right)* Selected images from a time-lapse sequence of an early embryo expressing endogenous ROD-1 tagged with GFP. GFP::ROD-1 is enriched in nuclei and localizes transiently to holocentric kinetochores in mitosis. In addition, GFP::ROD-1 starts to forms filaments during mitosis at the 8-cell stage, but not earlier (see also Movie S1). Dashed lines mark cell boundaries. Scale bar, 5 *µ*m. Blow-up, 2 *µ*m. **(B)** Selected images from a time-lapse sequence documenting the formation of GFP::ROD-1 filaments during mitosis at the 8-cell stage (see also Movie S2). mCherry::histone H2B labels chromosomes. Filaments, typically several *µ*m in length, form in the nucleus before NEBD and segregate to daughter cells by clustering at spindle poles. Kinetochore-localized GFP::ROD-1 is also visible (arrows). Time point 0 refers to the last frame before the appearance of GFP::ROD-1 on filaments and kinetochores. Scale bar, 2 *µ*m. **(C)** *(left)* Schematic highlighting the increase in nuclear GFP::ROD-1 levels during early embryonic development. *(right)* Selected images of nuclei from a time-lapse sequence of a developing embryo expressing GFP::ROD-1 that was followed from the 2-cell stage to the 8-cell stage. *(top)* Images show the EMS cell in the 4-cell embryo, which gives rise to the MS cell in the 8-cell embryo. *(bottom)* Likewise, the ABp cell gives rise to the ABpl cell. In both instances, nuclear GFP::ROD-1 levels increase gradually during the cell cycle, and are significantly higher in daughter cells. Time point 0 denotes the last frame before GFP::ROD-1 appears on kinetochores (EMS, ABp) or filaments (MS, ABpl). Similar results were obtained for nuclei of the P lineage (not shown). Scale bar, 2 *µ*m. **(D)** Quantification of average GFP::ROD-1 signal in nuclei and the cytoplasm in developing embryos as shown in *(C)*. Average fluorescence intensity was determined in images acquired every 20 s, averaged for the indicated number *n* of cells from at least 8 embryos, and plotted against time. Time point 0 denotes the last frame before the appearance of GFP::ROD-1 on filaments and/or kinetochores. Values are shown as mean ± 95% confidence interval for nuclear signal and as the mean for cytoplasmic signal. **(E)** *(left)* Quantification of nuclear GFP::ROD-1 levels in cells at different developmental stages. Measurements correspond to the last frame before GFP::ROD-1 appears on filaments and/or kinetochores (time point 0 s), showing a significant increase of nuclear signal at the 8-cell stage. Mild *rod-1(RNAi)* was used to reduce GFP::ROD-1 levels. Values are shown as mean ± 95% confidence interval. Statistical significance was determined by one-way ANOVA followed by Bonferroni’s multiple comparison test. \*\*\*\**P* < 0.0001; \*\**P* < 0.01. *(right)* Table showing that filament formation commences strictly at the 8-cell stage and can be completely suppressed by mildly reducing GFP::ROD-1 levels. **(F)** Selected images from 8-cell embryos whose AB lineage cells are going through mitosis. Mild depletion of GFP::ROD-1 slightly lowers enrichment in nuclei, which suppresses filament formation but does not prevent GFP::ROD-1 localization to kinetochores. Scale bar, 5 *µ*m.

We sought to investigate why GFP::ROD-1 filaments formed at this specific time in development. Quantification of the GFP::ROD-1 signal revealed that nuclear levels of GFP::ROD-1 gradually increased with every embryonic division, as well as during the course of each cell cycle, reaching a peak before NEBD (Fig. 4C - E). Consequently, nuclear GFP::ROD-1 levels before NEBD in 8-cell embryos were significantly higher than nuclear levels in all previous divisions (Fig. 4C - E). This suggested a straightforward explanation for the invariant timing of filament formation, namely that GFP::ROD-1 must become sufficiently concentrated before oligomerization is triggered. To directly test this idea, we used a mild RNAi regime to slightly reduce total GFP::ROD-1 levels in the embryo (Fig. 4E, F). In these RNAi conditions, nuclear GFP::ROD-1 levels in 8-cell embryos were comparable to those in control embryos at the 2-cell stage (Fig. 4E). Strikingly, GFP::ROD-1 no longer formed filaments in 8-cell embryos or at later developmental stages, despite the still prominent enrichment in nuclei and normal localization to mitotic kinetochores (Fig. 4E, F). These results demonstrate that ROD-1 has a propensity to oligomerize into filaments and suggest that oligomerization can be triggered locally by concentrating ROD-1 above a critical threshold.

### The ROD-1 β-propeller suppresses ubiquitous and complete oligomerization of ROD-1 into filaments

In human cells, deletion of the ROD β-propeller prevents kinetochore expansion, which may at least in part be a consequence of Spindly’s inability to bind RZZ. Because oligomerization of *C. elegans* ROD-1 into filaments was independent of SPDL-1^Spindly^, we wanted to address the role of the ROD-1 β-propeller. We therefore generated animals expressing mCherry::ROD-1 without β-propeller (Δ1-372) from an RNAi-resistant transgene integrated in single copy at a defined chromosomal locus (Fig. 5A). We found that mCherry::ROD-1(Δ1-372), just like full-length mCherry::ROD-1, localized to early embryonic nuclei and mitotic kinetochores when endogenous ROD-1 was present (Fig. 5B). By contrast, when we depleted endogenous ROD-1 by RNAi, mCherry::ROD-1(Δ1-372), but not full-length mCherry::ROD-1, oligomerized into filaments measuring up to 15 *µ*m in length that were ubiquitously present throughout the cytoplasm of the oocyte-producing gonad and early embryo, irrespective of developmental and cell cycle stage (Fig. 5B, D; Fig. S5A). Oligomerization of mCherry::ROD-1(Δ1-372) into filaments appeared to be essentially complete judging by the lack of diffuse cytoplasmic signal (Fig. 5B). Consequently, after depletion of endogenous ROD-1, mCherry::ROD-1(Δ1-372) was no longer detectable at mitotic kinetochores, and, as predicted by RZZ’s essential role at kinetochores, dividing embryos exhibited chromosome bridges in anaphase and were inviable (Fig. 5B, C). We conclude that ROD-1’s propensity to oligomerize into filaments is antagonized by its N-terminal β-propeller. The behavior of *C. elegans* ROD-1(Δ1-372) was unexpected given that human ROD(Δ1-375) prevented kinetochore expansion. While the molecular basis for this difference is currently unclear, the experiments in human cells and *C. elegans* identify the Rod β-propeller as a key regulator of RZZ self-assembly. The ROD-1(Δ1-372) mutant also illustrates that cells must have tight regulatory mechanisms in place to control Rod’s tendency to oligomerize.

**Figure 5:**
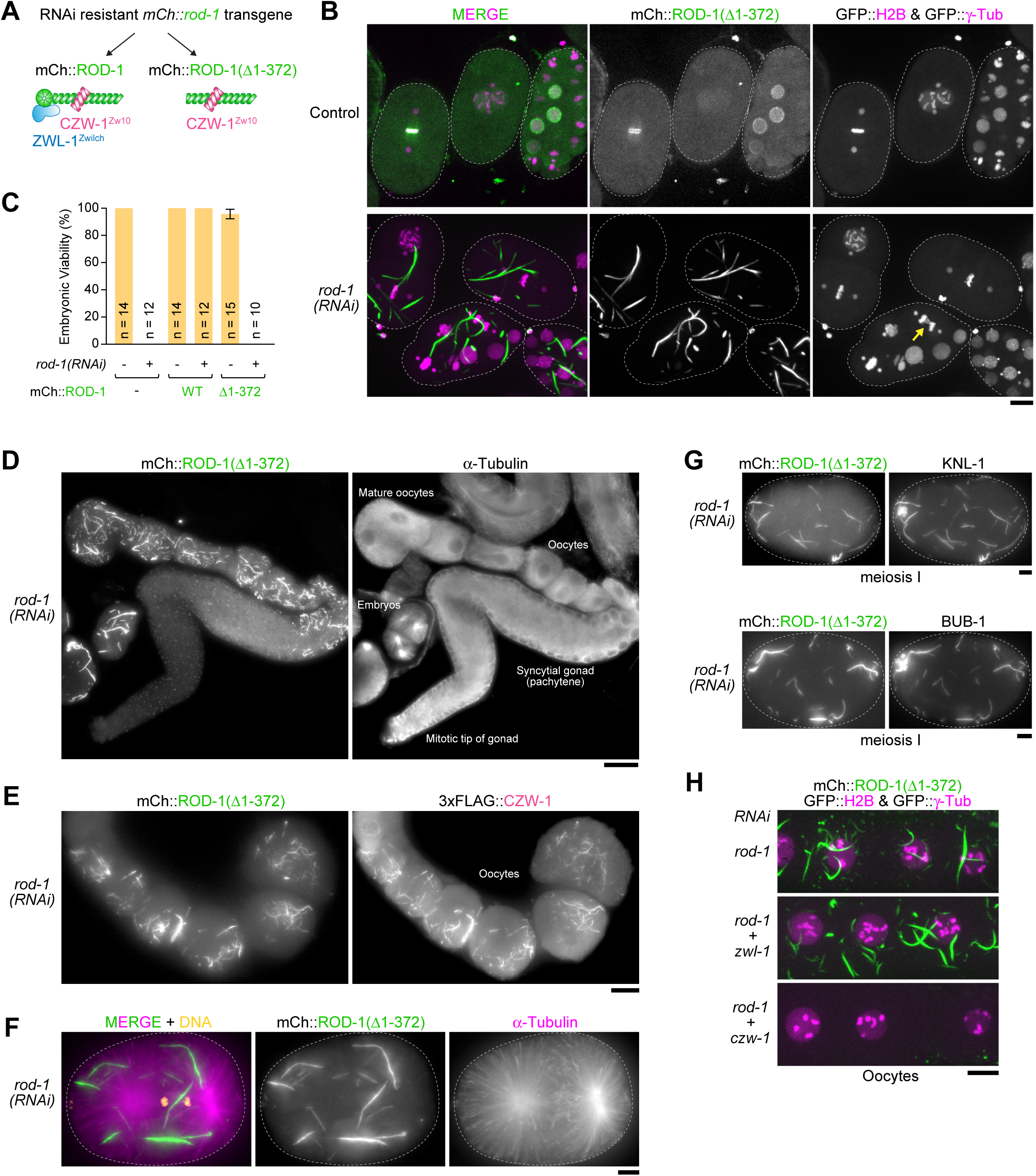
The ROD-1 β-propeller suppresses ubiquitous and complete oligomerization of ROD-1 into filaments. **(A)** Schematic of the protein complexes generated by the expression of RNAi-resistant *mCherry::rod-1* transgenes integrated in single copy on chromosome II. Note that the actual RZZ complex is a dimer of the trimer depicted here, which means mCherry::ROD-1(Δ1-372) is able to dimerize with endogenous ROD-1. **(B)** Selected images from time-lapse sequences in early embryos co-expressing mCherry::ROD-1(Δ1-372), GFP::histone H2B, and GFP::γ-tubulin. mCherry::ROD-1(Δ1-372) oligomerizes into large filaments when endogenous ROD-1 is depleted (see also Movie S3). This causes defects in chromosome segregation (arrow), because RZZ is titrated away from kinetochores. Dashed lines outline the embryos. Scale bar, 10 *µ*m. **(C)** Embryonic lethality assay showing that embryos expressing mCherry::ROD-1(Δ1-372) are only viable when endogenous ROD-1 is present. Values are plotted as mean ± 95% confidence interval, and *n* indicates the number of mothers whose progeny was counted. **(D)** Immunofluorescence image of an isolated oocyte-producing gonad, showing that mCherry::ROD-1(Δ1-372) forms filaments ubiquitously in oocytes and embryos. Scale bar, 25 *µ*m. **(E)** Immunofluorescence image of maturing oocytes showing co-localization of mCherry::ROD-1(Δ1-372) with 3xFLAG-tagged CZW-1^Zw10^ on filaments. Scale bar, 10 *µ*m. **(F)** Immunofluorescence image showing that mCherry::ROD-1(Δ1-372) filaments do not cluster at spindle poles during mitosis. Dashed lines outline the embryos. Scale bar, 5 *µ*m. **(G)** Immunofluorescence images showing co-localization of KNL-1 and BUB-1 with mCherry::ROD-1(Δ1-372) filaments in meiosis I embryos. Dashed lines outline the embryos. Scale bars, 5 *µ*m. **(H)** Fluorescence images of live oocytes showing that the formation of mCherry::ROD-1(Δ1-372) filaments depends on CZW-1^Zw10^ but not ZWL-1^Zwilch^. Scale bar, 10 *µ*m.

Immunostaining confirmed that endogenously tagged 3xFLAG::CZW-1^Zw10^ co-localized with mCherry::ROD-1(Δ1-372) on filaments (Fig. 5E), while ZWL-1^Zwilch^ and SPDL-1^Spindly^ were not detectable (Fig. S5B and data not shown). This agrees with prior work showing that CZW-1^Zw10^ binds to ROD-1’s Sec39 domain, which is present in ROD-1(Δ1-372), and that the ROD-1 β-propeller binds to ZWL-1^Zwilch^ and SPDL-1^Spindly^ [15,32]. Unlike the GFP::ROD-1 filaments (Fig. 4B; Fig. S4), mCherry::ROD-1(Δ1-372) filaments never clustered at mitotic spindle poles and showed no evidence of directed movement (Fig. 5F), consistent with the lack of SPDL-1^Spindly^ on these filaments. In the −1 oocyte and meiosis I embryo, where small ROD-1-dependent filaments are naturally present, the significantly larger filaments of mCherry::ROD-1(Δ1-372) were prominently decorated with KNL-1 and BUB-1 (Fig. 5G; Fig. S5C). KNL-3, NDC-80, and CENP-C also localized to the filaments, but to a lesser extent (data not shown). The association of kinetochore proteins with mCherry::ROD-1(Δ1-372) filaments was strictly limited to the −1 oocyte and meiosis I embryo (Fig. S5C), implying that their recruitment is regulated by posttranslational modifications and/or as-yet-unknown meiosis-specific factors. These results demonstrate that ROD-1(Δ1-372) and CZW-1^Zw10^ retain the ability to interact with other kinetochore proteins after assembly into higher-order oligomers.

### A complex of ROD-1(Δ1-372) and CZW-1^Zw10^ self-assembles efficiently into higher-order oligomers *in vitro*

We next sought to exploit the unique behavior of the ROD-1(Δ1-372) mutant to dissect the molecular requirements for ROD-1 oligomerization. Depletion of CZW-1^Zw10^ by RNAi completely suppressed filament formation of mCherry::ROD-1(Δ1-372) (Fig. 5H), likely because CZW-1^Zw10^ is essential for ROD-1 stability [15]. By contrast, mCherry::ROD-1(Δ1-372) filaments were not affected by depletion of ZWL-1^Zwilch^ or SPDL-1^Spindly^, consistent with the observation that ZWL-1^Zwilch^ and SPDL-1^Spindly^ did not localize to filaments. Furthermore, mCherry::ROD-1(Δ1-372) filaments readily formed after depletion of KNL-1, BUB-1, the Mis12 complex subunit KNL-3, and NDC-80, as well as after depletion of the mitotic regulators ICP-1^INCENP^, CDK-1, and PLK-1 (data not shown). This indicated that a complex of ROD-1(Δ1-372) and CZW-1^Zw10^ may be both necessary and sufficient for efficient oligomerization into filaments. To directly test this idea, we expressed the full-length RZZ complex and the ROD-1(Δ1-372)-CZW-1^Zw10^ complex in insect Sf21 cells using baculovirus (Fig. 6A). For purification and visualization, ROD-1 was N-terminally tagged with 6xHis followed by GFP, and CZW-1^Zw10^ contained an N-terminal StrepTagII. After a first affinity purification step with nickel resin, size exclusion chromatography (SEC) showed that GFP::RZZ fractionated as a complex with equal stoichiometry of its three subunits and without other obvious protein contaminants (Fig. 6A). After SEC, we examined the peak fractions by fluorescence microscopy but found no evidence that GFP::ROD-1 was present in filaments (Fig. 6B). We then applied the same purification procedure to the GFP::ROD-1(Δ1-372)-CZW-1^Zw10^ complex, which we obtained in similar amounts and purity as GFP::RZZ (Fig. 6A). In contrast to GFP::RZZ, GFP::ROD-1(Δ1-372)-CZW-1^Zw10^ fractionated in a broader peak in SEC. Subsequent examination of peak fractions using fluorescence and transmission electron microscopy revealed that GFP::ROD-1(Δ1-372) was present in filaments that were typically several *µ*m in length (up to ∼15 *µ*m) and had an invariant width of ∼50 nm (Fig. 6B - G). Filaments frequently associated laterally with each other to form loose bundles (Fig. 6C, F, G). Staining for the StrepTagII confirmed that CZW-1^Zw10^ co-localized with GFP::ROD-1(Δ1-372) on filaments (Fig. 6D). We also readily obtained filaments with a ROD-1(Δ1-372)-CZW-1^Zw10^ complex, confirming that the GFP tag was dispensable for ROD-1 oligomerization (Fig. S6). Once formed, filaments remained static and were stable at 4°C for several weeks, but did not tolerate freezing or high salt (500 mM NaCl), suggesting oligomerization involves electrostatic interactions. If the SEC step was omitted and partially purified GFP::ROD-1(Δ1-372)-CZW-1^Zw10^ was examined directly after nickel affinity chromatography, filament formation could occasionally be followed by live imaging, which revealed that filaments grow from both ends (Fig. 6E). We conclude that a purified ROD-1(Δ1-372)-CZW-1^Zw10^ complex oligomerizes efficiently into *µ*m-scale filaments *in vitro*, and that these filaments resemble those observed with this ROD-1 mutant *in vivo*. Thus, ROD-1(Δ1-372)-CZW-1^Zw10^ represents a minimal RZZ construct sufficient for self-assembly into higher-order oligomers.

**Figure 6:**
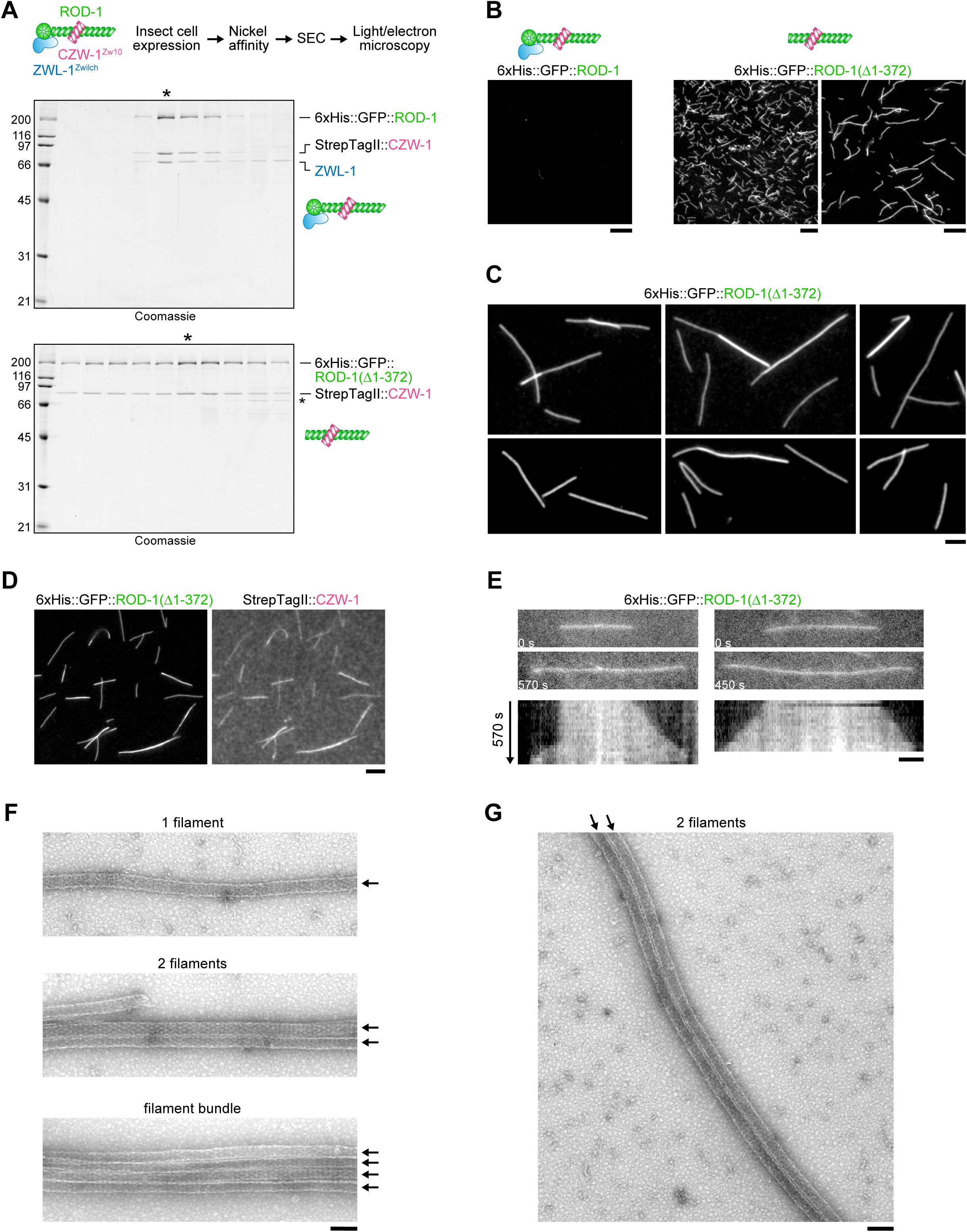
A complex of ROD-1 without β-propeller and CZW-1^Zw10^ self-assembles efficiently into higher-order oligomers *in vitro*. **(A)** *(top)* Workflow used to generate the *in vitro* data shown in this figure. *(bottom)* Coomassie-stained gels showing the protein fractions after SEC. Fractions analyzed in subsequent panels are marked with an asterisk. Note that purity and amount is comparable for of both complexes. Molecular weight is indicated in kDa on the left. **(B)** Images of the protein fractions marked with an asterisk in *(A)* examined by fluorescence light microscopy, showing that GFP::ROD-1(Δ1-372) but not full-length GFP::ROD-1 oligomerizes into *µ*m-long filaments with high efficiency. Scale bars, 10 *µ*m for left and middle image; 5 *µ*m for right image. **(C)** Higher magnification views of GFP::ROD-1(Δ1-372) filaments showing evidence of lateral bundling. Note that filaments reach up to 15 *µ*m in length. Scale bar, 2 *µ*m. **(D)** Fluorescence image confirming that GFP::ROD-1(Δ1-372) co-localizes with StrepTagII::CZW-1^Zw10^ on filaments *in vitro*. **(E)** Selected images *(top)* and corresponding kymographs *(bottom)* from a time-lapse sequence (30 s between frames), showing that GFP::ROD-1(Δ1-372) filaments grow from both ends. In this experiment, filaments were directly examined after the Nickel affinity step. Scale bar, 2 *µ*m. **(F), (G)** Transmission electron microscopy images of GFP::ROD-1(Δ1-372) filaments, showing that they have an invariant diameter of ∼50 nm and tend to associate laterally. Scale bars, 100 nm.

## DISCUSSION

The RZZ complex is well-established as an outer kinetochore component required for the recruitment of Mad1-Mad2 and dynein [11,33–37]. Here, we present evidence that RZZ additionally functions as a structural precursor for the assembly of the dynamic outermost kinetochore domain, which expands in early prometaphase to accelerate spindle assembly [22]. Recent work has revealed that the RZZ subunit Rod is evolutionarily related to coat proteins that form higher order assemblies on membranes [16,29]. We demonstrate in *C. elegans* that Rod proteins are also capable of self-assembly. Although additional components may be required *in vivo*, we show *in vitro* that a complex consisting of ROD-1 without its β-propeller and CZW-1^Zw10^ is sufficient to form *µ*m-scale filaments, which have an invariant diameter of ∼50 nm. Based on the human RZZ structure [16], the precursor of these filaments likely corresponds to a ROD-1(Δ1-372)-CZW-1^Zw10^ dimer. Similarly, the precursor of the GFP::RZZ filaments we observe *in vivo* is likely a dimer of a trimer, containing two copies each of ROD-1, CZW-1^Zw10^, and ZWL-1^Zwilch^. Deletion of the β-propeller greatly favors oligomerization of ROD-1 *in vitro* and *in vivo*. The ROD-1 β-propeller forms a tight complex with ZWL-1^Zwilch^ [15,32], raising the possibility that the presence of ZWL-1^Zwilch^ in RZZ has an inhibitory effect on self-assembly. However, depletion of ZWL-1^Zwilch^ in animals expressing mCherry::ROD-1 does not result in the ubiquitous formation of filaments we observe with mCherry::ROD-1(Δ1-372) [15]. This argues that the inhibitory effect of the ROD-1 β-propeller on RZZ self-assembly is direct. The extent to which CZW-1^Zw10^ contributes to self-assembly is difficult to assess, because CZW-1^Zw10^ is essential for the stability of ROD-1 [15]. Future work may reveal the spatial arrangement of precursors within ROD-1(Δ1-372)-CZW-1^Zw10^ filaments and provide insight into the self-assembly mechanism.

Based on our analysis of *C. elegans* RZZ, we suggest that human ROD’s central role in kinetochore expansion is directly related to its capacity for oligomerization. We envision that filamentous RZZ assemblies form the structural scaffold of the expanded kinetochore domain and function as dynamic recruitment platforms for corona proteins, including SAC components, the motors dynein and CENP-E, and CENP-F. In contrast to the KMN network, protein-protein interactions among components in the expanded outer domain are low-affinity and therefore challenging to dissect with biochemical approaches. Our finding that the expanded domain can be fully detached from the underlying kinetochore, either by acute CDK1 inhibition or by combining GFP::ROD over-expression with Knl1 depletion, suggests that RZZ-dependent expansion results in a distinct multi-protein assembly that is at least transiently stable without the KMN network. Consistent with recent studies [13,17], our findings imply that kinetochores contain two distinct pools of Mad1-Mad2: one pool recruited by the KMN network through a direct interaction with Bub1 [7–9], and another pool localized to the expanded outer domain, most likely recruited through a direct interaction with RZZ. The presence of BUBR1 and BUB3 on detached crescents may indicate that the expanded domain is able to generate MCC independently of BUB1.

Given RZZ’s propensity to form oligomeric assemblies, an important question is how this behavior is spatially restricted to kinetochores. Our observations in *C. elegans* embryos expressing endogenous GFP::RZZ show that local precursor concentration is a critical parameter governing oligomerization behavior. Specifically, analysis of GFP::RZZ filament formation suggests that the complex is refractory to oligomerization until a threshold density of precursors is achieved. This may in part explain why we failed to observe filaments with GFP::RZZ *in vitro*, as the concentrations we obtained in the preparations were relatively low. In the context of kinetochores, recruitment of RZZ by Knl-1-Bub1 could bring about the spatial proximity of RZZ precursors necessary to trigger oligomerization, which then proceeds autonomously. An important implication of the mechanism we propose is that self-recruitment contributes significantly to overall RZZ levels at unattached kinetochores. Thus, even low levels of Knl1-Bub1 may support kinetochore expansion, so long as a minimal number of RZZ precursors can be recruited to start filament assembly. This idea is consistent with the observation that kinetochores are able to expand in GFP::ROD-expressing cells depleted of KNL1. While Knl1-Bub1 appear to be dispensable for expansion *per se*, our results in GFP::ROD-expressing cells show that Knl1-Bub1 are required for the physical linkage between the expanded domain and the underlying kinetochore.

We also show that RZZ’s direct binding partner Spindly makes an important contribution to kinetochore expansion. This function of Spindly is separable from its function as the kinetochore adaptor for dynein-dynactin [15], because DCTN1 depletion shows that dynein-dynactin are dispensable for kinetochore expansion. Previous work indicates that Spindly is recruited to kinetochores through a direct interaction with the ROD β-propeller [15,16], and, consistently, deletion of the ROD β-propeller suppresses kinetochore expansion [15]. However, the ZWILCH(E422A/D426A) mutant shows that Spindly recruitment by RZZ *per se* is not sufficient for kinetochore expansion. Work in *C. elegans* suggested that SPDL-1^Spindly^, in addition to binding the ROD-1 β-propeller, also makes contact with ZWL-1^Zwilch^ [15]. The interaction between SPDL-1^Spindly^ and ZWL-1^Zwilch^ involves the conserved acidic residues we mutated in human ZWILCH, raising the possibility that an interaction between Spindly and ZWILCH may play role in RZZ self-assembly. Mitotic kinases have been implicated in the regulation of kinetochore expansion in *Xenopus* egg extracts [21]. It is therefore likely that kinetochore-localized kinase activity acts on RZZ and/or Spindly to promote self-assembly of RZZ. Mps1 is a particularly attractive candidate given that a recent study identified multiple Mps1-dependent phospho-sites on both Spindly and the ROD β-propeller [38].

In summary, we propose that recruitment of RZZ to kinetochores by Knl1-Bub1 locally concentrates Rod to favor self-assembly. Aided by Spindly and possibly kinase activity, this generates RZZ-based filaments that engage weakly-associating corona proteins to form the supramolecular assemblies visible as crescents and fibrous coronas by light and electron microscopy, respectively. Disassembly of the expanded domain after end-on microtubule attachment is likely achieved by dynein-dependent poleward transport of corona proteins, a process that is well documented [11,39–42]. Indeed, failure to completely disassemble the corona may contribute to the partial retention of MAD1 at microtubule-attached kinetochores that is observed in human Spindly mutants deficient in dynein recruitment [43,44]. Thus, RZZ not only helps build the expanded kinetochore but also recruits the motor that brings about its eventual compaction.

## MATERIALS AND METHODS

### Cell lines

cDNA coding for residues 1-591 of wild-type ZWILCH and the ZWILCH mutant E422A/D426A were cloned into a pcDNA5/FRT/TO-based vector (Invitrogen) modified to contain N-terminal Myc-GFP-TEV-S-peptide. Point mutations were subsequently introduced to make the transgene resistant to siRNA. Stable isogenic HeLa Flp-In T-REx cell lines expressing ZWILCH were generated by FRT/Flp-mediated recombination, as described previously [15].

### *C. elegans* strains

Worm strains (Table S1) were maintained at 20°C on standard NGM plates seeded with OP50 bacteria. To generate a strain stably expressing mCherry::ROD-1(373-2177) from chromosome II (locus ttTi5605), we used the same Mos1 transposon-based approach described previously for the strain expressing mCherry::ROD-1(1-2177) [15]. Integration of the transgene was verified by PCR and sequencing. Endogenous GFP tagging of the *rod-1* locus was done using CRISPR/Cas9-mediated genome editing, as described [45]. The repair template for *gfp::rod-1* included *gfp(S65C)* with introns, inserted upstream of the *rod-1* start codon via a GGRAGS linker, and the homology arms (1046 bp left; 1036 bp right). The PAM site for the guide RNA (5’-CCACAGCTTTTGCTTCGCCT-3’) was mutated from AGG to AGA in the repair template. To modify the *rod-1* locus using Cas9-triggered homologous recombination, the repair template (50 ng/*µ*L) was co-injected with two separate plasmids, one expressing guide RNA under the U6 promoter (50 ng/*µ*L) and the other expressing Cas9 under the eft-3 promoter (30 ng/*µ*L) into N2 worms [46]. The injection mix also contained three plasmids encoding fluorescent markers [pCFJ90 (P*myo-2::mCherry*, 2.5 ng/*µ*L), pCFJ104 (P*myo-3::mCherry*, 5 ng/*µ*L) and pGH8 (P*rab*-*3::mCherry*, 10 ng/*µ*L) that allowed selection for F1 transgenic animals. Progeny of F1 transgenic animals were screened for integration of *gfp* by PCR using a primer within *gfp* and a primer outside the left homology arm. Strains were outcrossed 6x with the wild-type N2 strain and other fluorescent markers were subsequently introduced by mating.

### RNA interference and drug treatments in HeLa cells

HeLa cells were maintained at 37 °C in a 5% CO_2_ atmosphere in Dulbecco’s modified Eagle’s medium (Gibco) supplemented with 10% fetal bovine serum (Gibco), 100 units/mL penicillin, 100 units/mL streptomycin, and 2 mM GlutaMAX (Gibco). For immunofluorescence, cells were seeded on 12-mm poly-L-lysine-coated coverslips in 12-well plates 24 h prior to transfection with siRNAs. Cells were transfected with siRNAs (Dharmacon On-Target plus; Table S2) targeting ROD/KNTC1, DCTN1, KNL1/CASC5, ZWILCH, and SPDL1/CCDC99, as described previously [43]. An siRNA against luciferase was used as a control. Transgene expression (Fig. 1 K - M; Fig. 2C; Fig. 3B - D; Fig. S2) was induced with 0.2 *µ*g/mL tetracycline (Sigma-Aldrich) 24 h post-transfection and cells were fixed 20 - 24 h later. To depolymerize microtubules prior to immunofluorescence, cells were treated with 1 *µ*M nocodazole (Sigma-Aldrich) for 4 h. To detach kinetochore crescents from centromeres (Fig. 2; Fig. S1), cells were incubated for 4 h with 1 *µ*M nocodazole followed by a 30-min incubation with nocodazole and 10 *µ*M CDK1 inhibitor RO-3306 (Sigma-Aldrich).

### RNA interference in *C. elegans*

For production of double stranded RNA (dsRNA; Table S3), oligos with tails containing T3 and T7 promoters were used to amplify regions from genomic N2 DNA (gDNA) or cDNA. PCR reactions were cleaned (NucleoSpin Gel and PCR Clean-up, Macherey-Nagel) and used as templates for T3 and T7 transcription reactions (MEGAscript, Invitrogen). Transcription reactions were cleaned (NucleoSpin RNA Clean-up, Macherey-Nagel) and annealed in soaking buffer (3x soaking buffer is 32.7 mM Na_2_HPO_4_, 16.5 mM KH_2_PO_4_, 6.3 mM NaCl, 14.1 mM NH_4_Cl). dsRNA was delivered by injecting L4 hermaphrodites, and animals were processed for live-imaging or immunofluorescence after incubation at 20°C for 45 - 50 h. For the experiments in Fig. 4E and F, a partial knockdown of ROD-1 in strain OD3367 was performed by injecting young adults and incubating injected animals at 20 °C for 5 h.

### Embryonic viability

Embryonic viability assays were performed at 20°C. L4 hermaphrodites injected with dsRNA were grown for 40 h on NGM plates containing OP50 bacteria, single adults were placed on new mating plates (NGM plates with a small amount of OP50 bacteria) and removed 8 h later. The number of hatched and unhatched embryos on each plate was counted after further incubation for 16 h.

### Indirect immunofluorescence

For immunofluorescence experiments in cultured human cells, HeLa cells were fixed immediately after aspiration of the medium with 4% paraformaldehyde in Phem buffer (60 mM Pipes, 25 mM Hepes, 10 mM EGTA, 2 mM MgCl_2_, pH 6.9) for 5 min at room temperature, then permeabilized for 2 min with 0.1% Triton X-100 in Phem buffer and rinsed once with Phem buffer. For stainings with anti-ZW10 antibody, cells were fixed in methanol at −20 °C for 45 min, and re-hydrated for 5 min in PBS / 0.5% Triton X-100 followed by 5 min in PBS / 0.1% Triton X-100. Cells were blocked for 30 min in AbDil solution (PBS, 4% IgG-free BSA [Jackson Immuno Research], 0.1% Triton X-100, 5 mM NaN_3_) and incubated with primary antibody over night at 4 °C, diluted in AbDil (mouse anti-CENP-E clone 1H12, 1:500 [Abcam ab5093]; sheep anti-CENP-F, 1:800 [gift from Stephen Taylor]; guinea pig anti-CENP-C, 1:1500 [MBL PD030]; rabbit anti-Spindly OD174, 1:5000 [gift from Arshad Desai]; goat anti-GFP, 1:15000 [gift from Anthony Hyman]; mouse anti-p150, 1:500 [BD Transduction Laboratories 610473]; mouse anti-MAD1 clone BB3-8, 1:300 [Millipore MABE867]; mouse anti-MAD2, 1:500 [Santa Cruz Biotechnology sc-65492]; rabbit anti-ZWILCH, 1:900 [gift from Andrea Musacchio]; rabbit anti-ZW10, 1:300 [gift from Andrea Musacchio]; human anti-centromere antibodies ACA, 1:5000 [Antibodies Incorporated]; sheep anti-BUBR1, 1:1000 [gift from Stephen Taylor]; sheep anti-BUB3, 1:250 [gift from Stephen Taylor]; mouse anti-BUB1, 1:200 [Abcam ab54893]; mouse anti-HEC1 clone 9G3, 1:1000 [Abcam ab3616]; rabbit anti-KNL1 OD111, 1:2000 [gift from Arshad Desai]; rabbit anti-DSN1 OD110, 1:1500 [gift from Arshad Desai]. After washing for 3×5 min in PBS / 0.1% Triton X-100, cells were incubated with secondary antibodies conjugated to fluorescent dyes (Alexa 488, 594, 647 [Jackson ImmunoResearch]). Cells were washed again for 3×5 min, rinsed in PBS and mounted in Prolong Gold with DAPI stain (Invitrogen).

Immunofluorescence in *C. elegans* embryos was carried out as described [47], using the following primary antibodies: goat anti-GFP, 1:15000; rabbit anti-mCherry OD78, 1 *µ*g/mL; mouse anti-α-Tubulin DM1a, 1:1000 (Sigma-Aldrich); mouse anti-FLAG M2, 1:1000 (Sigma-Aldrich); rabbit anti-KNL-1 OD33, 1 *µ*g/mL; rabbit anti-BUB-1 OD31, 1 *µ*g/mL; rabbit anti-ZWL-1 OD85, 1 *µ*g/mL; rabbit anti-SPDL-1 OD164, 1:7000. Antibodies OD31, OD33, OD78, OD85, and OD164 were a gift from Arshad Desai.

Images were recorded on a Zeiss Axio Observer microscope controlled by ZEN 2.3 software, using a 100x NA 1.46 Plan-Apochromat objective, an Orca Flash 4.0 camera (Hamamatsu), and an HXP 200C Illuminator (Zeiss). Images of HeLa cells were processed using the deconvolution module for ZEN 2.3.

### Live-imaging of *C. elegans* embryos

Adult gravid hermaphrodite worms were dissected in a watch glass filled with Egg Salts medium (118 mM KCl, 3.4 mM MgCl_2_, 3.4 mM CaCl_2_, 5 mM HEPES, pH 7.4), and embryos were mounted on a fresh 2% agarose pad and covered with an 18 mm×18 mm coverslip (No. 1.5H, Marienfeld). Imaging was performed in a temperature-controlled room at 20°C using a Nikon Eclipse Ti microscope coupled to an Andor Revolution XD spinning disk confocal system, composed of an iXon Ultra 897 CCD camera (Andor Technology), a solid-state laser combiner (ALC-UVP 350i, Andor Technology), and a CSU-X1 confocal scanner (Yokogawa Electric Corporation), controlled by Andor IQ3 software (Andor Technology).

### Imaging of purified 6xHis::GFP::ROD-1(Δ1-372)-StrepTagII::CZW-1

Fractions containing purified protein complex were analyzed by putting a drop of 4 *µ*L on an acid washed 13-mm diameter coverslip and inverting the coverslip onto a slide. For visualization of StrepTagII::CZW-1, protein fractions were first incubated for 10 min with Strep-Tactin conjugated to Oyster 645 (1:300, IBA). Images were recorded on a Zeiss Axio Observer microscope (system as described for indirect immunofluorescence) using a 40x NA 1.3 or 63x NA 1.4 Plan-Apochromat objective. For time-lapse imaging (Fig. 6E), a 5×1 *µ*m z-stack was recorded every 30 s.

### Transmission electron microscopy

Protein fractions were allowed to settle on 300 mesh Formvar-carbon-coated nickel grids and were negatively stained with a 2% aqueous solution of uranyl acetate. Grids were examined with a JEM1400 transmission electron microscope (JEOL) operating at 120 kV. Images were acquired using a post-column high-resolution (11 megapixels) high-speed camera (SC1000 Orius, Gatan).

### Image analysis

Image analysis was performed using Fiji software (Image J version 2.0.0-rc-56/1.51 h).

#### Quantification of kinetochore volumes and fluorescence intensity in HeLa cells

Volume and intensity of kinetochore fluorescence for CENP-E, CENP-F, Spindly, and GFP were measured using the 3D Objects Counter tool. To each image stack (0.2-*µ*m z-steps) encompassing one cell, the Subtract Background function was applied (rolling ball radius of 5 pixels), and a mask was generated based on an empirically determined threshold that maximized the number of detected kinetochores, while minimizing the detection of false objects. The mask was re-directed to the original, unprocessed image stack to determine the number of voxels and the integrated intensity for each object, which could consist of one or several closely apposed kinetochores. Object values were then summed to give the total number of voxels and the total integrated intensity for kinetochores in the image stack. To determine background intensity, the number of voxels and integrated intensity were measured for five separate regions in the image stack that did not contain kinetochores. Values for the five regions were averaged, normalized to the total number of kinetochore voxels, and subtracted from the total integrated kinetochore intensity. The total number of kinetochores in the image stack was determined separately using the CENP-C signal after maximum intensity projection of the image stack, followed by thresholding, automatic particle counting, and verification by visual inspection. Finally, the total number of kinetochore voxels and the total integrated kinetochore intensity were divided by the total number of kinetochores to yield the average number of voxels and integrated intensity per kinetochore per cell. Average kinetochore volumes and integrated intensities were determined for 20 cells in two independent experiments.

#### Quantification of GFP::ROD-1 intensity in C. elegans embryos

Measurements of nuclear and cytoplasmic GFP::ROD-1 signal (Fig. 4D, E) were performed after maximum intensity projection of 12×1 *µ*m z-stacks, captured every 20 s in embryos developing from the 2-cell to the 8-cell stage. The mean intensity in the nucleus or in the cytoplasm of the same cell was determined, and the mean intensity of a region outside the embryos (camera background) was subtracted.

### Statistical analysis

Values in figures are reported as mean ± 95% confidence interval. Statistical analysis was performed with GraphPad Prism 7.0 software. The type of statistical analysis (two-tailed t-test or one-way ANOVA/Bonferroni’s multiple comparison test) is indicated in the figure legends. Differences were considered significant at *P* values below 0.05.

### Biochemistry

#### Expression constructs

The cDNAs coding for ZWL-1, CZW-1, and ROD-1 (residues 1-2177 and 373-2177) were cloned into the pACEbac1 expression vector. ROD-1 constructs were tagged N-terminally with 6xHis followed by GFP. CZW-1 contained an N-terminal StrepTagII. Bacmid recombination and virus production were carried out as described previously [48].

#### Purification of RZZ and R(Δ1-372)Z from insect cells

6xHis::GFP::ROD-1, ZWL-1 and StrepTagII::CZW-1 (RZZ) or 6xHis::GFP::ROD-1(Δ1-372) and StrepTagII::CZW-1 [R(Δ1-372)Z] were co-expressed in 500-mL cultures (SFM4 medium, Hyclone) of Sf21 cells (0.8×10^6^ cells/mL), co-infected with corresponding viruses. Cells were harvested by centrifugation at 800×g for 5 min. Pellets were re-suspended in lysis buffer [50 mM Hepes, 200 mM NaCl, pH 8.0] supplemented with EDTA-free complete Protease Inhibitor Cocktail (Roche), sonicated, and cleared by centrifugation at 34000×g for 40 min. Both complexes were purified by batch affinity chromatography using HIS-Select Nickel Affinity Gel beads (Sigma). Beads were washed with wash buffer [50 mM Hepes, 200 mM NaCl, 10 mM Imidazole, pH 8.0] and eluted on a gravity column with elution buffer [50 mM Hepes, 200 mM NaCl, 250 mM Imidazole, pH 8.0]. Protein complexes were further purified by size exclusion chromatography using a Superose 6 10/300 column (GE Health Care) equilibrated with 25 mM Hepes, 150 mM NaCl, pH 7.5. Glycerol and DTT were added to fractions containing RZZ or R(Δ1-372)Z to a final concentration of 10% (v/v) and 1 mM, respectively, and aliquots were kept at 4°C, as filaments did not tolerate freezing.

## ACKNOWLEDGEMENTS

The authors wish to thank Rui Fernandes and the Histology and Electron Microscopy Service at i3S for support and Helder Maiato for critical reading of the manuscript. Funding for this project was provided by the European Research Council under the European Union’s Seventh Framework Programme (ERC grant agreement n ERC-2013-StG-338410-DYNEINOME), by the European Molecular Biology Organization (EMBO Installation Grant 2545), by the Fundação para a Ciência e a Tecnologia (IF/01015/2013/CP1157/CT0006 to R.G. and SFRH/BPD/95648/2013 to C.P.), and by ‘Norte-01-0145-FEDER-000029 - Advancing cancer research: from basic knowledge to application’, supported by the Norte Portugal Regional Operational Programme (NORTE 2020), under the PORTUGAL 2020 Partnership Agreement through the European Regional Development Fund (FEDER).

## AUTHOR CONTRIBUTIONS

R. G., C. P., R. M. R., J. B. G., and A. X. C. conceived and designed experiments. C. P. performed the *C. elegans* work and the experiments with purified proteins. R. M. R. performed the work in cultured human cells. J. B. G. generated the baculovirus reagents and contributed to insect cell expression and protein purification. D. K. C. generated the *C. elegans* strain expressing GFP::ROD-1. R. G., C. P., and R. M. R. prepared the figures and wrote the manuscript with advice from A. X. C.

## FIGURE LEGENDS

**Figure S1:**
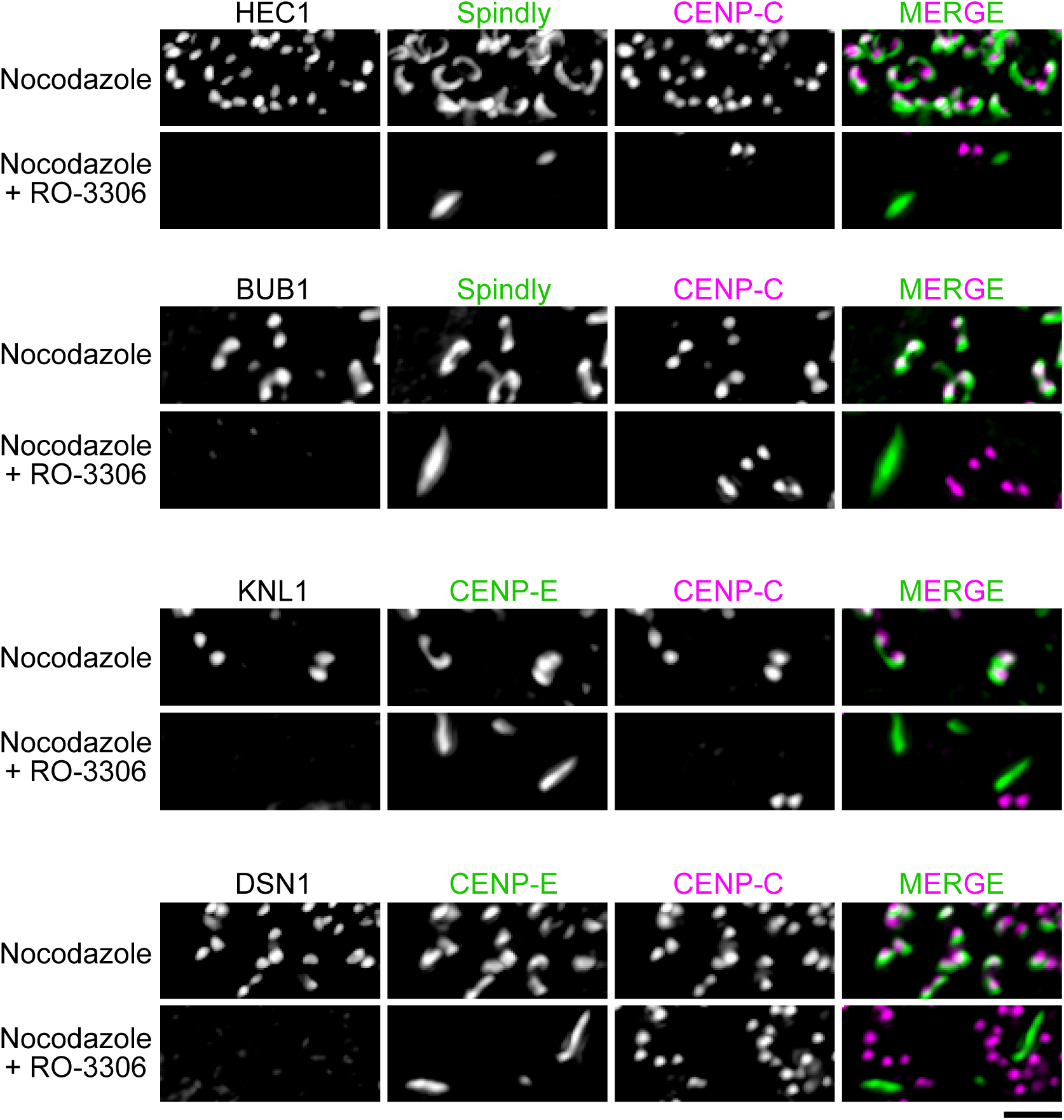
KMN network components do not localize to expanded kinetochore domains detached from centromeres after acute CDK1 inhibition. Immunofluorescence images showing co-localization of HEC1, BUB1, KNL1, and DSN1 with CENP-C in nocodazole-treated cells as a positive control, and the lack of signal on detached expanded kinetochore domains, marked by CENP-E or Spindly. Scale bar, 2 *µ*m.

**Figure S2:**
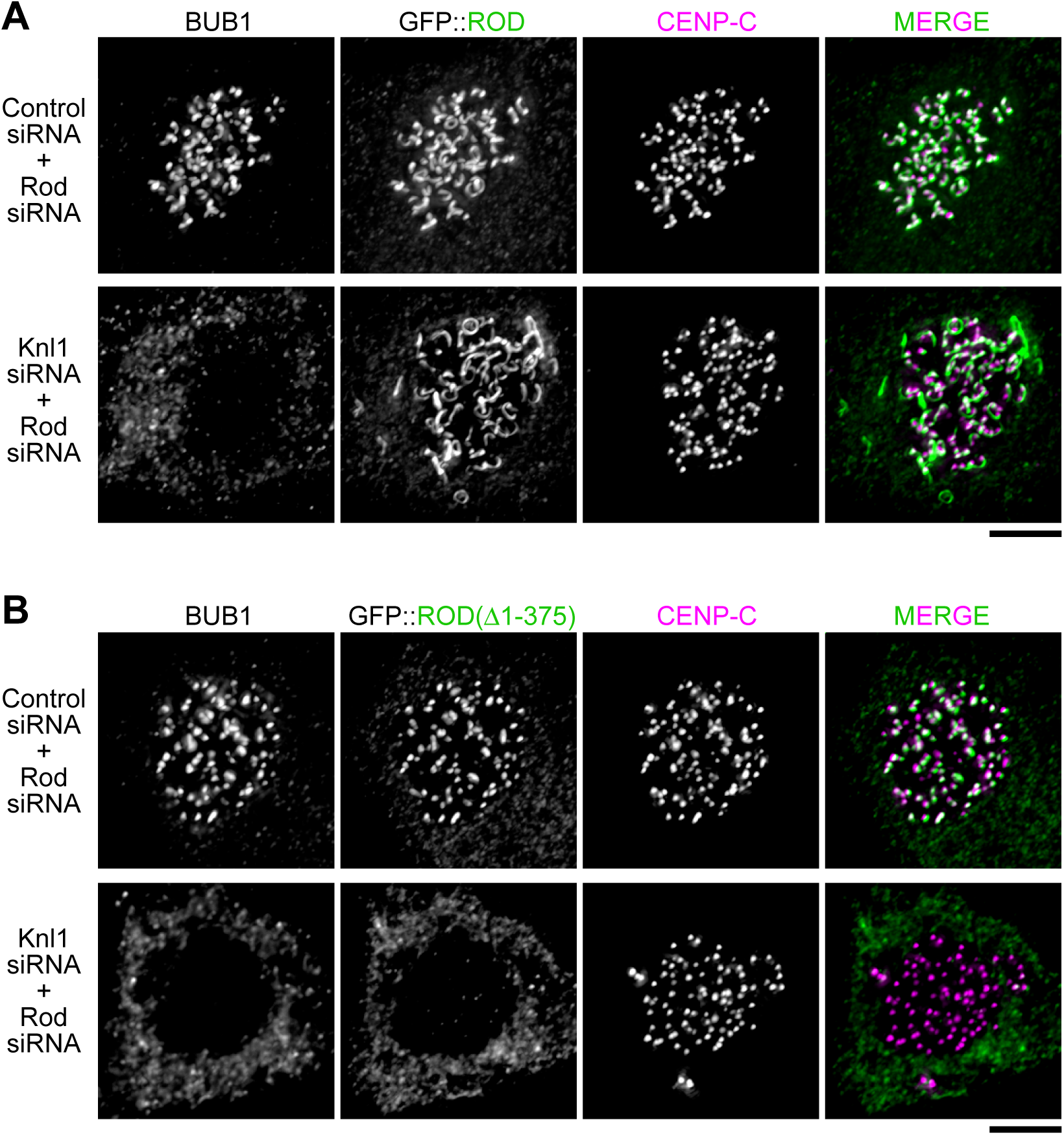
Kinetochore expansion promoted by exogenous GFP::ROD after depletion of KNL1 requires the ROD β-propeller. **(A)** Immunofluorescence images showing that RNAi-resistant GFP::ROD supports robust kinetochore expansion after depletion of endogenous ROD even when KNL1 is co-depleted. **(B)** Immunofluorescence images showing that GFP::ROD(Δ1-375) does not expand in nocodazole when endogenous ROD is depleted and cannot support expansion after co-depletion of KNL1. Scale bars, 5 *µ*m.

**Figure S3:**
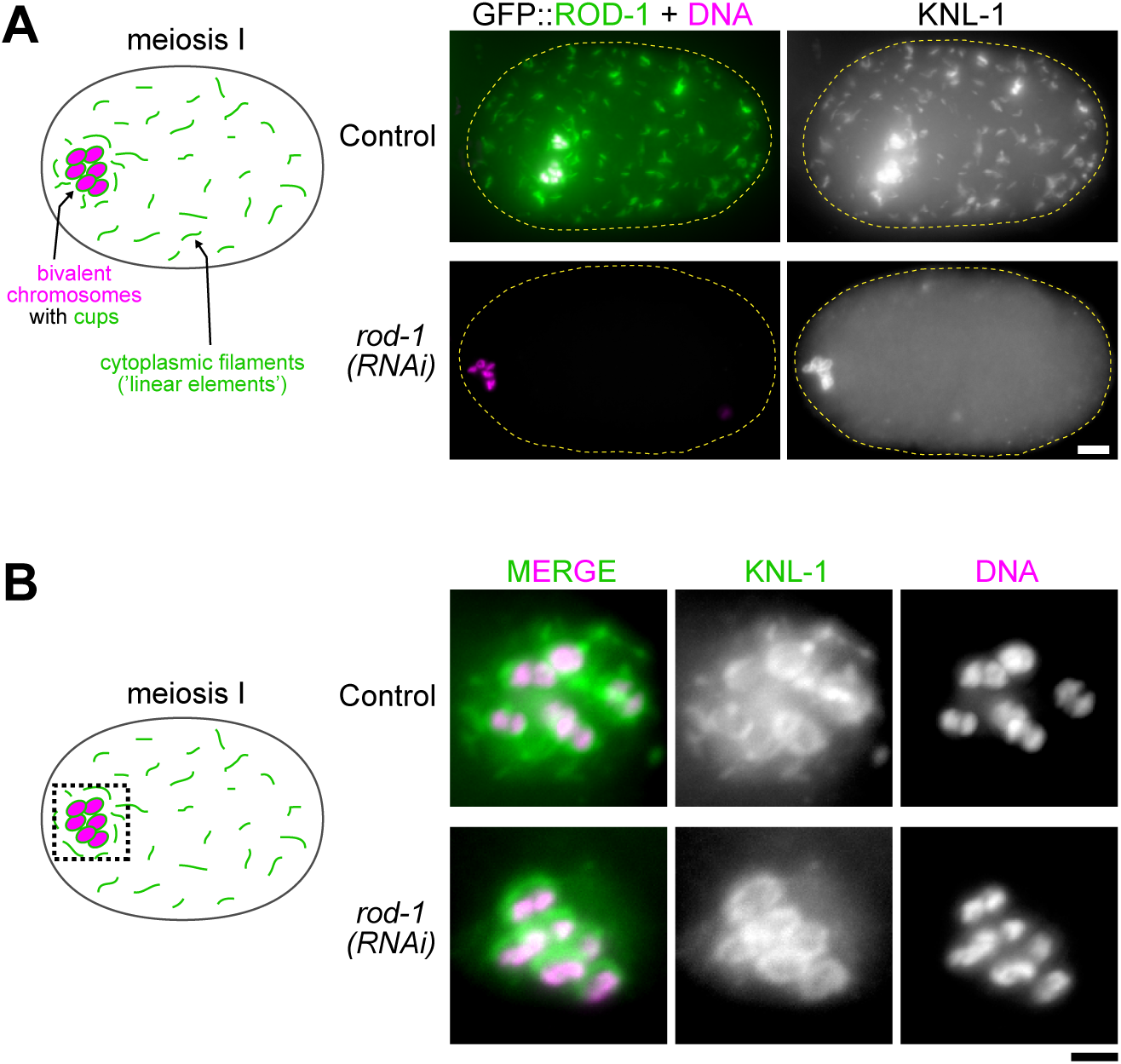
Cytoplasmic filaments containing KNL-1 are naturally present in the meiosis I embryo and depend on ROD-1. **(A)**, **(B)** Immunofluorescence images of the *C. elegans* meiosis I embryo, showing that GFP::ROD-1 and KNL-1 localize to small cytoplasmic filaments (linear elements) that are enriched around the bivalent chromosomes, as well as to cup-like meiotic kinetochores encircling the bivalents, as described previously [27,28]. Depletion of ROD-1 inhibits formation of linear elements but does not affect KNL-1 localization to meiotic kinetochores. Scale bars, 5 *µ*m *(A)* and 2 *µ*m *(B)*.

**Figure S4:**
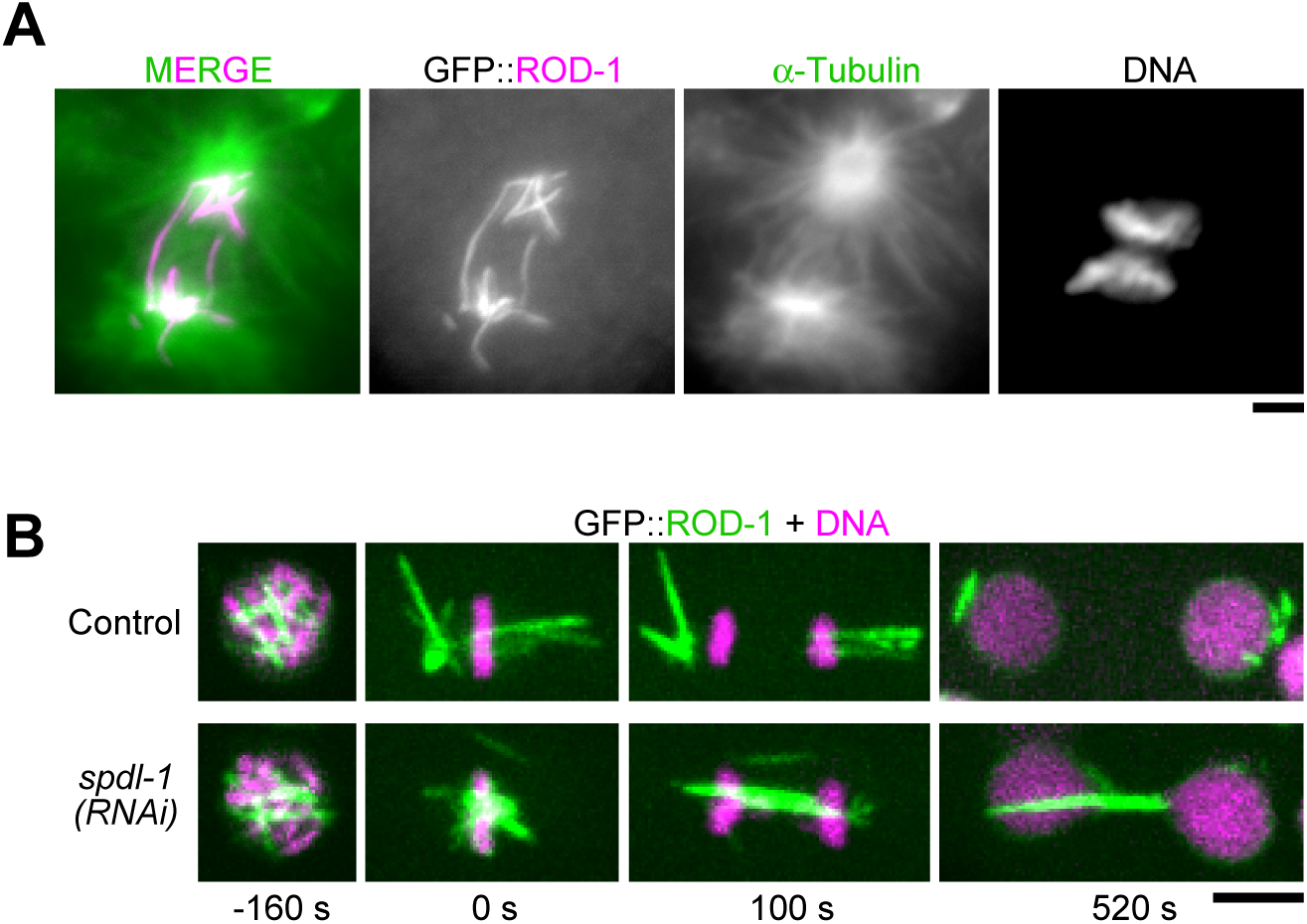
GFP::ROD-1 filaments segregate to daughter cells by clustering at spindle poles in a SPDL-1^Spindly^-dependent manner and disassemble upon mitotic exit. **(A)** Immunofluorescence image of a dividing cell in a multicellular embryo showing that GFP::ROD-1 filaments cluster at spindle poles. Scale bar, 2 *µ*m. **(B)** Selected images from a time-lapse sequence showing that clustering of GFP::ROD-1 filaments at spindle poles depends on SPDL-1^Spindly^. Images also show that GFP:ROD-1 filaments that cluster at spindle poles disassemble at the end of mitosis (time point 520 s). Scale bar, 5 *µ*m.

**Figure S5:**
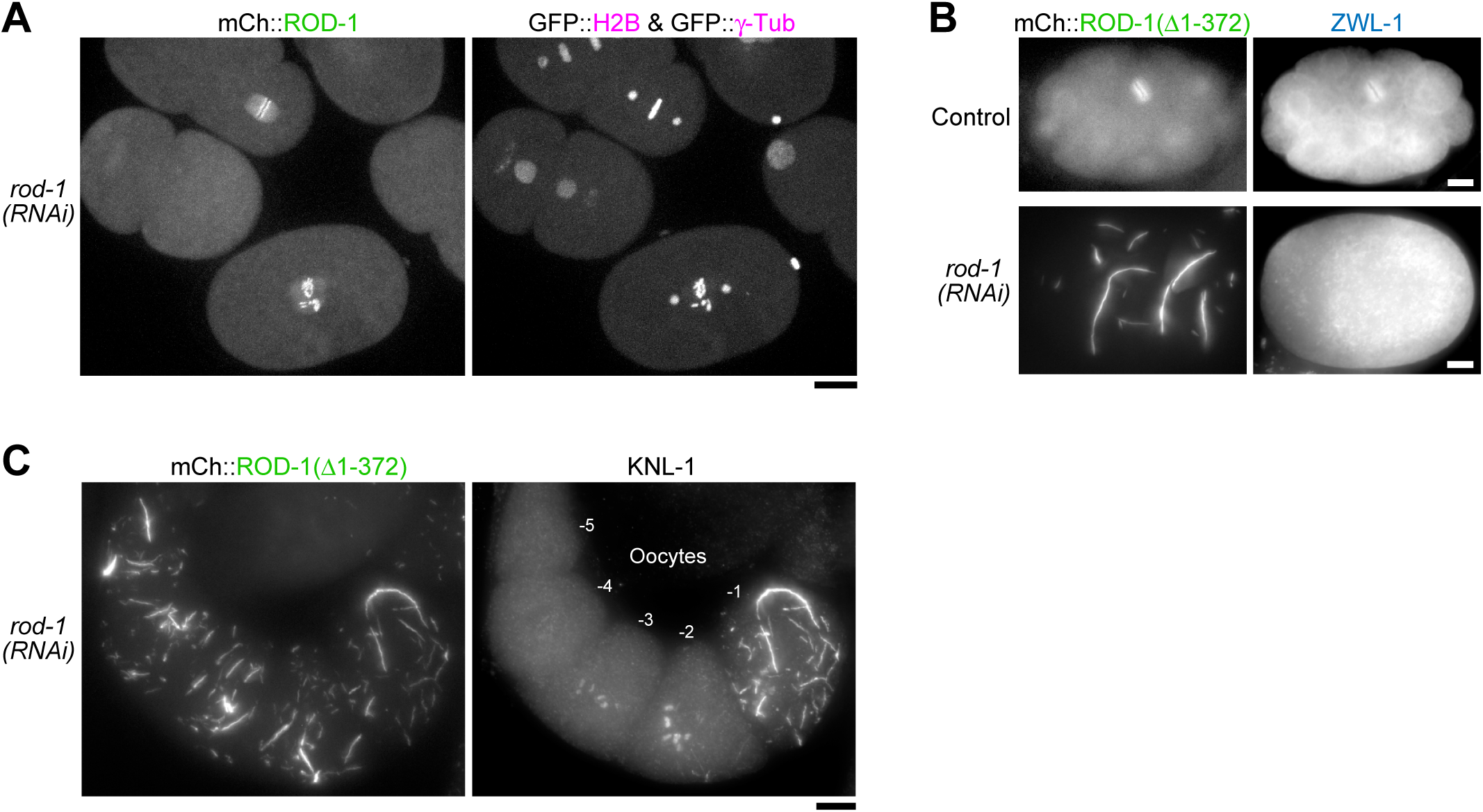
Additional analysis of filaments generated by mCherry::ROD-1(Δ1-372). **(A)** Selected image from a time-lapse sequences showing 1- and 2-cell embryos co-expressing full-length mCherry::ROD-1, GFP::histone H2B, and GFP::γ-tubulin. Depletion of endogenous ROD-1 has no effect on the localization of mCherry::ROD-1 to kinetochores, nor does it induce filament formation. Scale bar, 10 *µ*m. **(B)** Immunofluorescence image of a multi-cellular embryo *(top)* showing ZWL-1 signal at mitotic kinetochores as a positive control for the antibody, and a lack of ZWL-1 signal on mCherry::ROD-1(Δ1-372) filaments in a meiotic embryo *(below)*. Scale bars, 5 *µ*m. **(C)** Immunofluorescence image of oocytes demonstrating that KNL-1 localizes to mCherry::ROD-1(Δ1-372) filaments specifically in the most mature −1 oocyte. Scale bar, 10 *µ*m.

**Figure S6:**
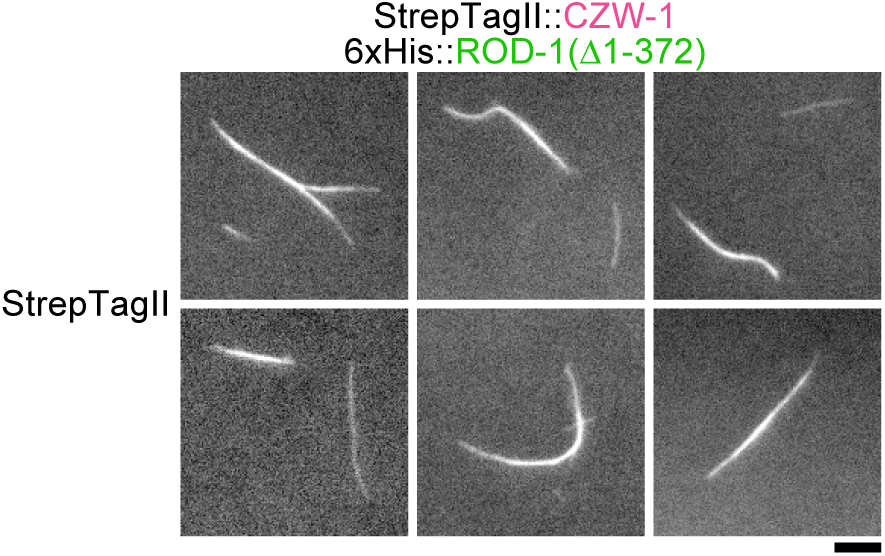
Self-assembly into filaments of ROD-1(Δ1-372)-CZW-1^Zw10^ *in vitro* does not require an N-terminal GFP tag on ROD-1(Δ1-372). Fluorescence images showing filaments assembled *in vitro* with purified 6xHis::ROD-1(Δ1-372)-StrepTagII::CZW-1^Zw10^. Filaments were detected with Strep-Tactin conjugated to Oyster 645. Scale bar, 1 *µ*m.

**Table S1:** C. elegans strains

| Strain Name | Genotype |
| --- | --- |
| N2 | *wild type (ancestral N2 Bristol)* |
| GCP24 | *ltSi426[pDC192; Prod-1::mCherry::rod-1::rod-1 3'UTR; cb-unc-119(+)] II; unc-119(ed3) III?; ruls32[pAZ132; Ppie-1::GFP::his-58; cb-unc-119(+)] III; ddls6[Ppie-1::GFP::tbg-1; cb-unc-119(+)] V \** |
| GCP287 | *prtSi103[pRG499; Prod-1::mCherry::rod-1(373-2177)::rod-1 3'UTR; cb-unc-119(+)] II; unc-119(ed3) III \** |
| GCP303 | *prtSi103[pRG499; Prod-1::mCherry::rod-1(373-2177)::rod-1 3'UTR; cb-unc-119(+)] II; unc-119(ed3) III?; ruls32[pAZ132; Ppie-1::GFP::his-58; cb-unc-119(+)] III; ddls6[Ppie-1::GFP::tbg-1; cb-unc-119(+)] V \** |
| GCP529 | *rod-1(lt62[gfp::rod-1]) IV; ltIs122[pAA64; Ppie-1::mCherry::his-58; cb-unc-119(+)]; unc-119(ed3) III? \** |
| GCP539 | *prtSi103[pRG499; Prod-1::mCherry::rod-1(373-2177)::rod-1 3'UTR; cb-unc-119(+)] II; unc-119(ed3) III?; czw-1[prt78(3xflag::czw-1)] IV \** |
| OD3367 | *rod-1(lt62[gfp::rod-1]) IV* |

\* *unc-119(ed3) III* was present in parental strains, but these strains have not been sequenced to determine whether the *unc-119* gene still contains the *ed3* mutation.

**Table S2:**
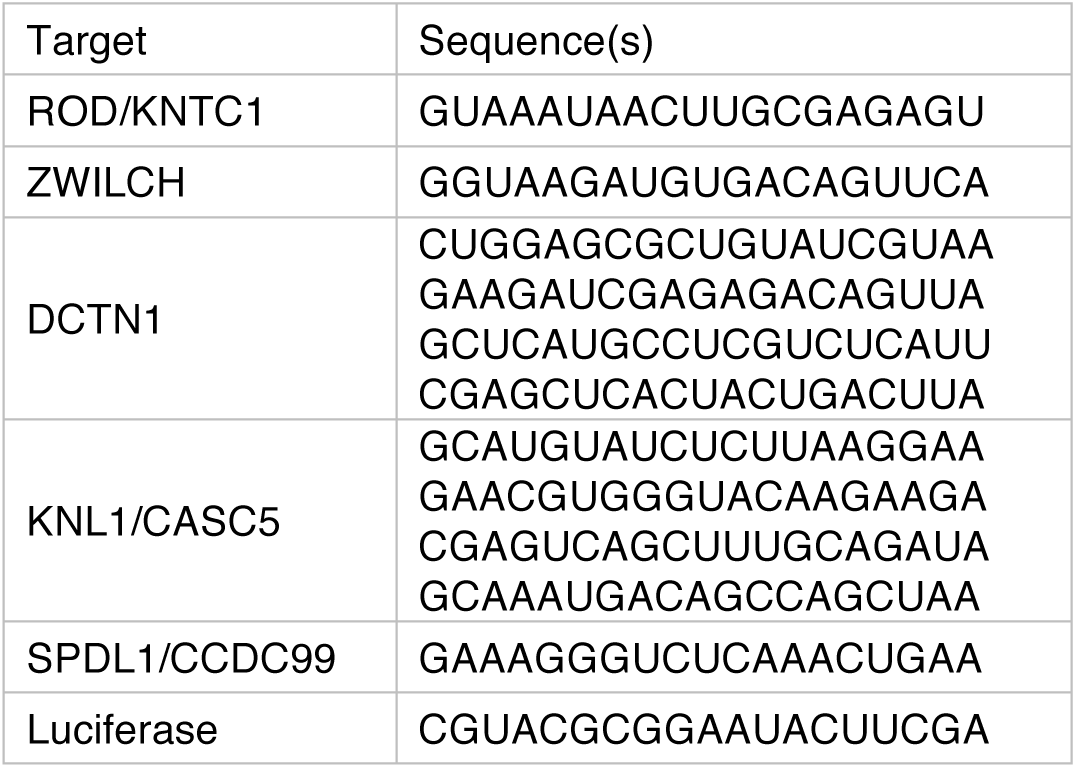
siRNAs

**Table S3:**
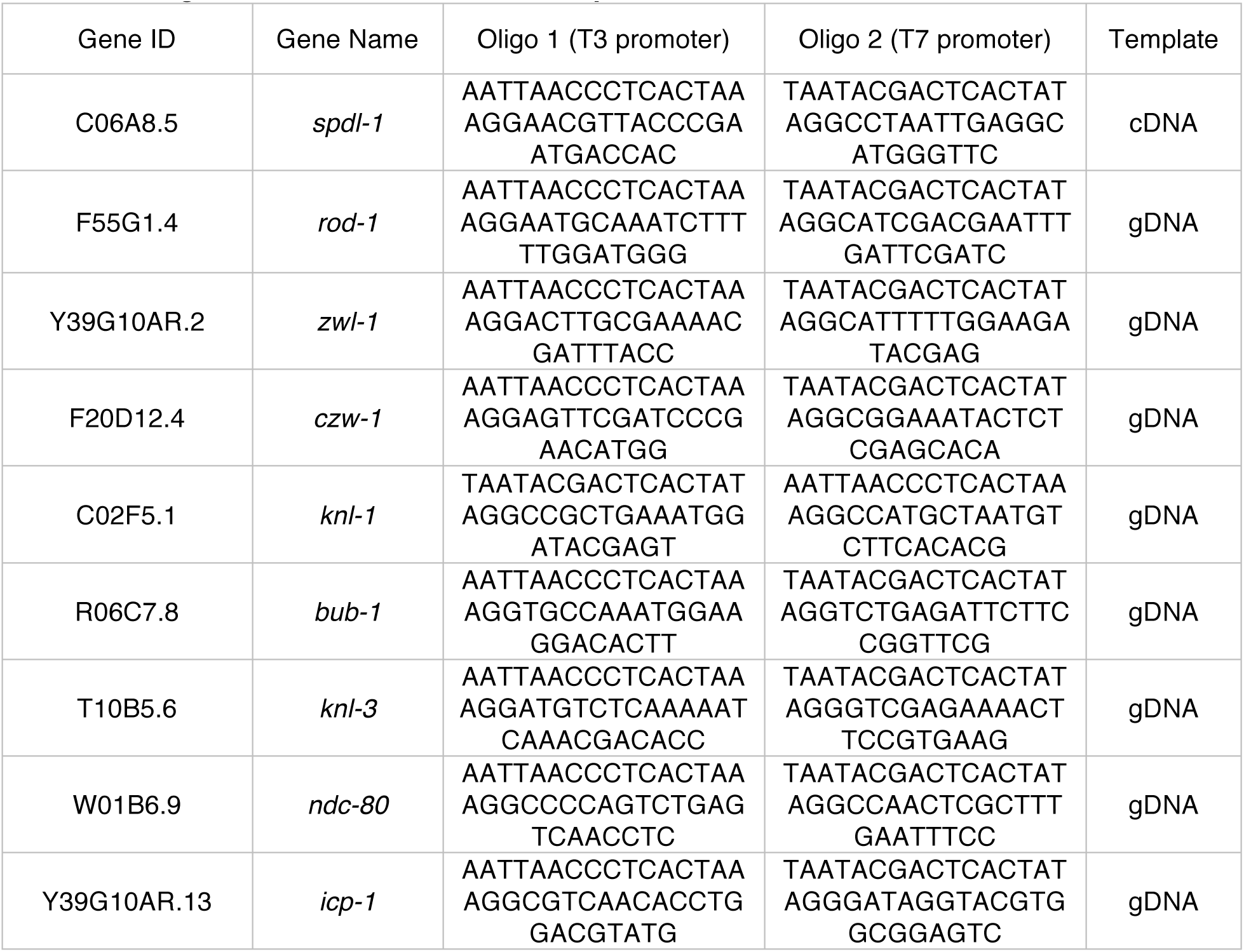
Oligos for double-stranded RNA production

**Movie S1:** GFP::ROD-1 localization during early embryonic development. Movie starts at anaphase in the 1-cell embryo and finishes at the 16-cell stage. Time lapse is 20 s and playback speed is 6 frames per second. Scale bar, 5 *µ*m.

**Movie S2:** AB cells going through mitosis in an 8-cell embryo co-expressing GFP::ROD-1 and mCherry::histone H2B. Time lapse is 20 s and playback speed is 6 frames per second. Scale bar, 5 *µ*m.

**Movie S3:** Early embryos co-expressing mCherry::ROD-1(373-2177), GFP::histone H2B, and GFP::γ-tubulin, after depletion of endogenous ROD-1 by RNAi. Time lapse is 10 s and playback speed is 6 frames per second. Scale bar, 10 *µ*m.

